# An axonemal intron splicing program sustains *Plasmodium* male development

**DOI:** 10.1101/2023.10.07.561333

**Authors:** Jiepeng Guan, Peijia Wu, Xiaoli Mo, Xiaolong Zhang, Wenqi Liang, Xiaoming Zhang, Lubing Jiang, Jian Li, Huiting Cui, Jing Yuan

**Affiliations:** State Key Laboratory of Cellular Stress Biology, Innovation Center for Cell Signal Network, School of Life Sciences, Xiamen University, Xiamen, Fujian 361102, China; Shanghai Institute of Immunity and Infection, Chinese Academy of Sciences, Shanghai, China

## Abstract

Differentiation of male gametocyte into flagellated fertile male gamete relies on the axoneme assembly, a major component of male development for mosquito transmission of malaria parasite. RNA-binding protein (RBP)-mediated post-transcription regulation plays important roles in eukaryotic sexual development, including the *Plasmodium* female development. However, the role of RBP in defining the *Plasmodium* male transcriptome and its function in the male gametogenesis remain elusive. Here, we screened the gender expression of the genome-wide RBPs and identified an undescribed male-specific RBP gene *Rbpm1* in the *Plasmodium*. RBPm1 is localized in the nucleus of male gametocytes. RBPm1-deficient parasites fail to assemble the axoneme for male gametogenesis and thus mosquito transmission. RBPm1 interacts with spliceosome E complex and regulates splicing initiation of certain introns in a group of 26 axonemal genes. RBPm1 deficiency results in intron retention and protein loss of these axonemal genes. Intron deletion restores axonemal proteins expression and partially rectifies axonemal defects in RBPm1-null gametocytes. Further splicing assays in both reporter and endogenous genes exhibit stringent recognition of the axonemal introns by RBPm1. Therefore, the splicing activator RBPm1 and its target introns constitute an axonemal intron splicing program in the post-transcription regulation essential for *Plasmodium* male development.

## Introduction

Malaria is a worldwide infectious disease caused by protozoan parasites *Plasmodium* [1]. The spread of *Plasmodium* depends on the transition between the mammal host and the *Anopheles* mosquito. In mammal hosts, a small proportion of intraerythrocytic asexual parasites undergo sexual development, irreversibly differentiating into the sexual precursor female and male gametocytes, which are transmission competent for the mosquito vector [2]. Within 10 min after being ingested into the mosquito midgut, the gametocytes develop into fertile gametes and escape from host erythrocytes, a process known as gametogenesis [3]. A flagellated motile male gamete fertilizes with a female one to form a zygote. After the zygote-ookinete-oocyst-sporozoite development in the mosquito, the parasites are finally injected from the salivary gland into a mammal host, completing the transmission of malaria parasite [4].

Sexual development plays a central role in malaria transmission [5, 6]. When activated by two joint mosquito stimuli (a temperature drop [7] and a metabolite xanthurenic acid [8]), a female gametocyte produces a haploid female gamete, while a male gametocyte gives rise to 8 haploid male gametes [3]. Female gametogenesis undergoes minor morphological changes, while male gametogenesis involves fast and spectacular changes [9, 10]. During the male gametogenesis, two spatially distinct components are coordinated. One is the cytoplasmic assembly of 8 basal bodies and axonemes, the other is the three successive rounds of genome replication without nuclear division, resulting in an octoploid nucleus. Subsequent release of 8 axonemes attached with chromosomes results in 8 flagellated daughter gametes from the cell body of male gametocytes, a process termed “exflagellation”.

Parasite stage transition during the *Plasmodium* life cycle requires a fine-tuned multilayer regulation of gene expression [11-13]. Previous studies have identified transcriptional and epigenetic programs critical for the sexual commitment and development of the gametocytes [14-23], however, how the *Plasmodium* establishes distinct repertoires of transcripts between the male and female gametocytes remains incompletely illustrated. RNA-binding protein (RBP) can interact with mRNA to regulate RNA metabolism and function, such as RNA processing, stability, localization, and translation [24]. It is well-known that the post-transcription regulation complex consisting of several RBPs, including the DOZI (development of zygote inhibited) and CITH (CARI/Trailer Hitch homolog), had been shown to repress the translation of multiple mRNAs in female gametocytes [25-27]. So far, our understanding of post-transcription control is limited in the male gametogenesis. Recent transcriptome studies in both human malaria parasite *P. falciparum* and mouse malaria parasite *P*. *berghei* revealed that certain RBPs are specifically or preferentially expressed in the male gametocytes [28, 29], implying gender-specific roles of RBP in the post-transcription regulation for male development. However, systematic identification of male RBPs for male gametogenesis and their precise roles in defining the gender distinct transcriptome via the post-transcription regulation have not been reported.

In this study, we performed the gametocyte gender transcriptome analysis and obtained a list of gender-specific RBPs in the rodent malaria parasite *P. yoelii*. From this list, we identified a functional unknown gene (PY17X_0716700, named *Rbpm1* in this study) showing greatest relative transcription at male compared to female gametocytes. We demonstrated that RBPm1 was a nuclear RBP essential in male gametogenesis and mosquito transmission of parasite. RBPm1 interacted with spliceosome E complex and initiated the splicing of certain introns in a group of axonemal genes. RBPm1-deficient parasites could not express the axonemal proteins and failed to assemble the axoneme. These findings revealed an RBPm1-mediated axonemal genes introns splicing program essential for *Plasmodium* male development.

## Results

### RNA-binding protein RBPm1 is expressed in nucleus of male gametocytes

Approximately 180 putative *Plasmodium* RBPs had been predicted *in silico* [30]. To identify the key RBPs for male gametogenesis, we searched the male gametocyte-specific RBPs in the rodent malaria parasite *P. yoelii*. Using the fluorescence-activated cell sorting, highly purified male and female gametocytes were collected from a *P. yoelii* reporter line *DFsc7* (Figure 1A and Fig S1A), in which fluorescent proteins GFP and mCherry are expressed in male and female gametocytes respectively [31]. We performed RNA-seq and obtained gender-specific gametocyte transcriptome data (Fig S1B). Among the 179 *P. yoelii* RBPs, an unstudied gene (PY17X_0716700) was identified with the greatest enrichment in transcription at male compared to female gametocytes (Figure 1B, left panel). This gene was named as *Rbpm1* for RBP in male gametocyte. Notably, the *Rbpm1* orthologs PBANKA_0716500 and PF3D7_0414500 are also the top male RBP genes of *P*. *berghei* and *P. falciparum* respectively (Figure 1B, middle and right panels) based on the gender gametocyte transcriptomes [28, 29].

**Figure 1.**
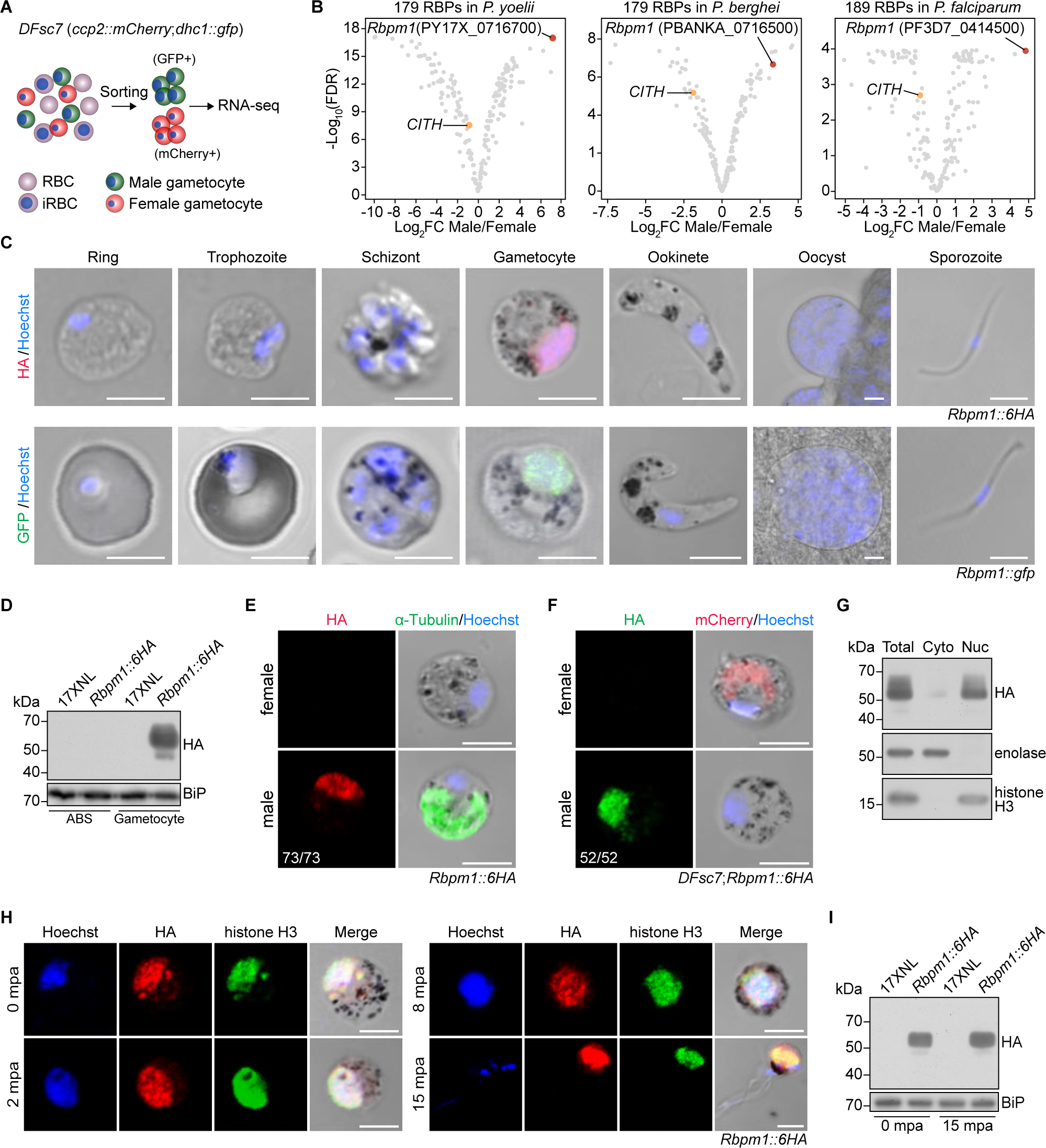
RBPm1 is a potential nuclear RBP in the male gametocytes. **A.** Flowchart showing the purification and transcriptome analysis of male (green, GFP+) and female (red, mCherry+) gametocytes from a *P. yoelii* parasite reporter line *DFsc7*. **B.** Gender analysis of gene transcription for the *Plasmodium* genome-wide putative RBPs between male and female gametocytes. The top male gene PY17X_0716700, RBPm1, is marked in red. CITH (orange dot) is a known female RBP. The results of *P. berghei* (middle panel) and *P. falciparum* (right panel) were based on the gametocyte transcriptome contributed by Yeoh, L.M. 2017 and Lasonder, E. 2016. **C.** Stage expression of RBPm1 during the *P. yoelii* life cycle. Immunofluorescence assay (IFA) of RBPm1 expression in the *Rbpm1::6HA* parasites stained with anti-HA antibody (top panel). Live cell imaging of the RBPm1::GFP protein in the *Rbpm1::gfp* parasites (bottom panel). Nuclei were stained with Hoechst 33342. Representative results from three independent experiments. Scale bars: 5 µm. **D.** Immunoblot of RBPm1 in the asexual blood stage (ABS) and gametocyte of the *Rbpm1::6HA* parasites. BiP as a loading control. Representative for three independent experiments. **E.** IFA of HA-tagged RBPm1 and α-Tubulin (male gametocyte marker protein) in *Rbpm1::6HA* gametocytes. Representative for three independent experiments. Scale bars: 5 µm. **F.** IFA of HA-tagged RBPm1 and mCherry (expressed in female gametocytes) in the *DFsc7*;*Rbpm1::6HA* gametocytes. Representative for three independent experiments. Scale bars: 5 µm. **G.** Immunoblot of RBPm1 in cytosolic and nuclear fractions of *Rbpm1::6HA* gametocytes. Enolase (cytoplasmic/Cyto) and histone H3 (nuclear/Nuc) proteins used as controls respectively. Representative for two independent experiments. **H.** IFA of HA-tagged RBPm1 and histone H3 during male gametogenesis of the *Rbpm1::6HA* parasites. mpa: minute post activation. Representative for three independent experiments. Scale bars: 5 µm. **I.** Immunoblot of RBPm1 expression in the *Rbpm1::6HA* parasites during male gametogenesis. Representative for two independent experiments.

To investigate RBPm1 expression during the parasite life cycle, we tagged endogenous RBPm1 with a sextuple HA (6HA) at the carboxyl (C)-terminus in the *P. yoelii* 17XNL strain (wildtype or WT) using CRISPR-Cas9 [32, 33]. The tagged parasite *Rbpm1*::*6HA* developed normally in mice and mosquitoes, indicating no effect of tagging on protein function. Immunofluorescent assay (IFA) showed that RBPm1 was expressed only at gametocytes, but not at asexual blood stages, ookinetes, oocysts, and sporozoites (Figure 1C, upper panel). Immunoblot also confirmed the gametocyte-restricted expression of RBPm1 (Figure 1D). Gametocyte-specific expression of RBPm1 was observed in another parasite line *Rbpm1*::*gfp* in which the RBPm1 was tagged with GFP from the 17XNL (Figure 1C, lower panel). To dissect whether RBPm1 is male-specific, the *Rbpm1*::*6HA* gametocytes were co-stained with antibodies against α-Tubulin (a male gametocyte marker) and HA. RBPm1 was only detectable in the male gametocytes (Figure 1E). Additionally, we tagged RBPm1 with 6HA in the reporter line *DFsc7* and observed the male specific expression of RBPm1 (Figure 1F). We noticed the nuclear localization of RBPm1 in all the male gametocytes tested (Figure 1C, E and F), which was further confirmed by immunoblot of nuclear and cytoplasmic fractions from the *Rbpm1*::*6HA* gametocytes (Figure 1G). Last, we analyzed the localization dynamics of RBPm1 throughout the entire process of gametogenesis (0, 2, 8, and 15 minutes post activation, mpa) in the *Rbpm1*::*6HA* parasites. Both IFA and immunoblot revealed consistent protein abundance and nuclear localization of RBPm1 during gametogenesis (Figure 1H and I). Together, these results demonstrated that RBPm1 was a nuclear protein specifically expressed in the male gametocytes.

### RBPm1 is essential for male gametogenesis and mosquito transmission of parasite

The *P. yoelii Rbpm1* gene encodes a protein of 361 amino acid (aa) residues, with two RNA recognition motifs (RRM1 and RRM2). To investigate its function, we generated a mutant line *ΔRbpm1* by deleting the entire genomic sequence (1904 bp) of *Rbpm1* gene in *P. yoelii* 17XNL strain using CRISPR-Cas9 (Figure 2A). *ΔRbpm1* produced normal level of male and female gametocytes in mice (Figure 2B), indicating that RBPm1 is not essential for asexual blood stage proliferation and gametocyte formation. We next measured the male gametogenesis by counting exflagellation centers (ECs) *in vitro* after stimulation with 50 μM xanthurenic acid (XA) at 22°C. *ΔRbpm1* showed a complete deficiency in EC formation (Figure 2C and D) and male gamete release (Figure 2E). In contrast, RBPm1 disruption had no impact on female gamete formation *in vitro* (Figure 2F), which corresponded with no RBPm1 expression in female. *ΔRbpm1* produced no ookinetes *in vitro* as well as midgut oocysts and salivary gland sporozoites in the infected mosquitoes (Figure 2G, H, and I), indicating transmission failure in mosquito. Additionally, we deleted each of the RNA recognition motifs RRM1 (119-190 aa) and RRM2 (203-274 aa) of endogenous RBPm1 in the 17XNL (Figure 2A). Both mutants, *Δrrm1* and *Δrrm2*, displayed similar defects as *ΔRbpm1* (Figure 2B and D), suggesting essential role of both two RNA recognition motifs in RBPm1 function.

**Figure 2.**
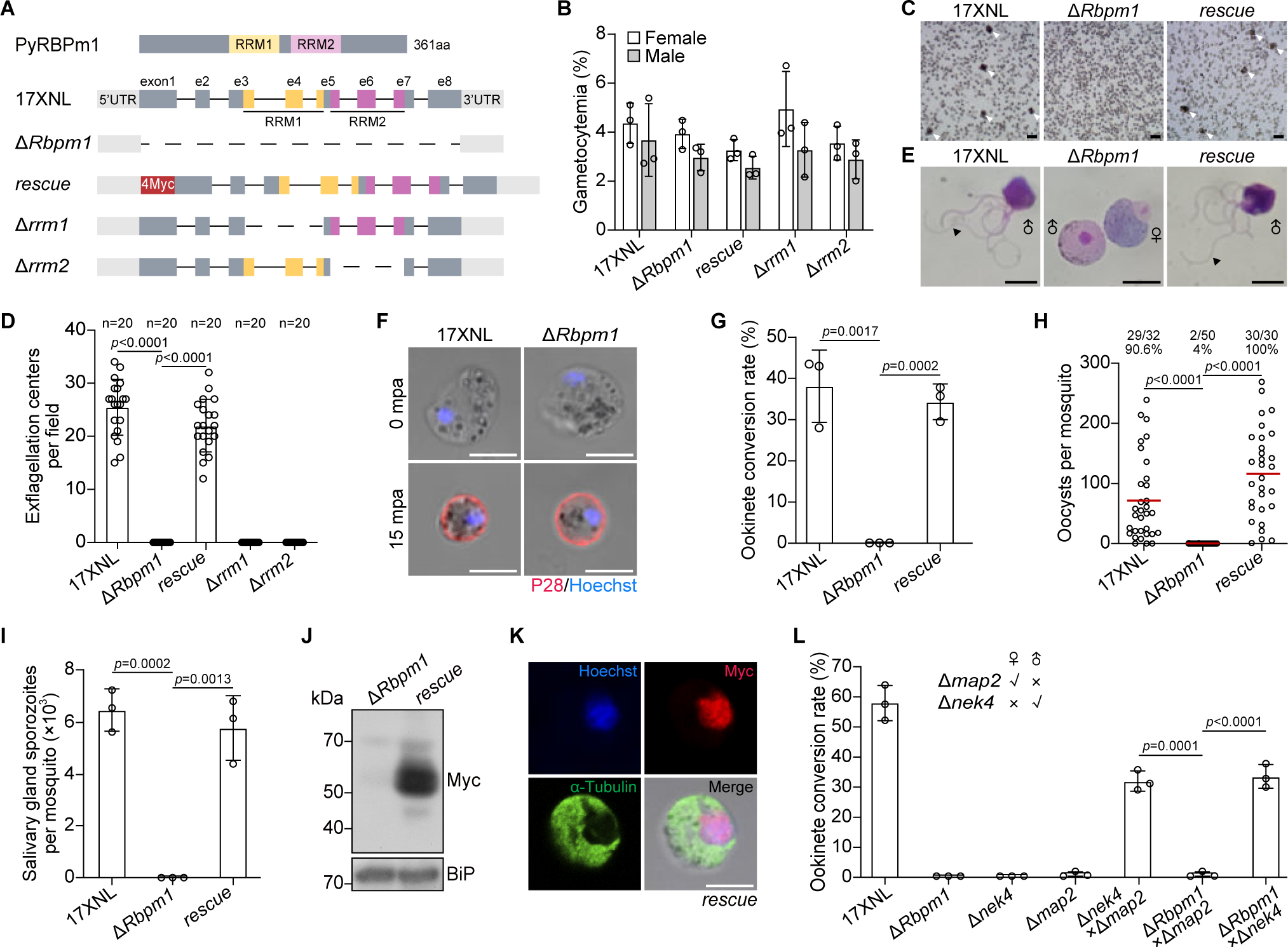
RBPm1 is essential for male gametogenesis and mosquito transmission. **A.** A schematic showing genetic modification at the *Rbpm1* locus in the *P. yoelii* parasite. The top panel depicts the protein structure of RBPm1 with two RNA recognition motifs RRM1 (residues 119-190, yellow) and RRM2 (residues 203-174, purple). Δ*Rbpm1*, deletion of the whole coding sequence from the 17XNL (wildtype) strain; *rescue*, the Δ*Rbpm1* line complemented with *Rbpm1* fused with a 4Myc tag; Δ*rrm1* and Δ*rrm2*, deletion each of RRM1 and RRM2 from the 17XNL. **B.** Female and male gametocyte formation in mice for the modified parasite. Data are means ±SEM of three independent experiments. **C.** Exflagellation center (EC) formation of activated male gametocytes at 10 mpa. Cell clusters representing the EC are marked with white arrows. Representative for four independent experiments. Scale bars: 20 μm. **D.** Quantification of EC formation. The ECs were counted within a 1×1-mm square area in the hemocytometer under a light microscope. n represents the number of fields counted. Means ± SEM, two-tailed *t*-test. Three independent experiments. **E.** Light microscope images (100×) of the exflagellated male gametes (black arrow) after Giemsa staining. Four independent experiments. Scale bars: 5 μm. **F.** Female gamete formation assayed by P28 staining. P28 is a female gamete plasma membrane protein. Three independent experiments. Scale bars: 5 μm. **G.** Ookinete formation *in vitro*. Data are means ± SEM from three independent experiments, two-tailed *t*-test. **H.** Midgut oocyst formation in mosquitoes at 7 days after blood feeding. x/y at the top represents the number of mosquitoes containing oocysts/the number of dissected mosquitoes, and the percentage represents the infection prevalence of mosquitoes. Red lines show the mean value of oocyst numbers, two-sided Mann–Whitney *U* test. Representative results from three independent experiments. **I.** Salivary gland sporozoite formation in mosquitoes at 14 days after blood feeding. At least 20 infected mosquitoes were counted in each group. Data are means ± SEM of three independent experiments, two-tailed *t*-test. **J.** Immunoblot analysis of RBPm1 expression in gametocytes of the complemented line *rescue*. BiP as a loading control. Two independent experiments. **K.** IFA of Myc-tagged RBPm1 and α-Tubulin in gametocytes of the *rescue* parasite. Representative for three independent experiments. Scale bars: 5 μm. **L.** Gender gamete fertility assay of the Δ*Rbpm1* by parasite genetic cross. Fertility was determined by ookinete development of Δ*Rbpm1* gametes after cross-fertilization with mutant lines that are defective in either female (Δ*nek4*) or male (Δ*map2*) gametes. Data are means ± SEM of three independent experiments, two-tailed *t*-test.

To further confirm that the *ΔRbpm1* phenotype was caused by *Rbpm1* deficiency, we introduced a sequence consisting of the coding region of *Rbpm1* and a N-terminal quadruple Myc epitope (4Myc) back to the *Rbpm1* locus in the *ΔRbpm1* line, generating the complemented line *rescue* (Figure 2A). The 4Myc-tagged RBPm1 was detected in the *rescue* gametocytes (Figure 2J) and localized at nucleus of male gametocytes (Figure 2K). The *rescue* parasites restored the formation of ECs (Figure 2C and D), male gametes (Figure 2E), ookinetes (Figure 2G), midgut oocysts (Figure 2H), and salivary gland sporozoites (Figure 2I).

Lastly, we performed genetic crosses between *ΔRbpm1* mutant and the male-deficient line Δ*map2* or the female-deficient line Δ*nek4*. As expected, the cross between Δ*map2* and Δ*nek4* produced the ookinetes *in vitro* (Figure 2L). The ookinete formation was restored in the Δ*Rbpm1* parasites that were crossed with Δ*nek4* but not Δ*map2*, further confirming the *ΔRbpm1* defects in male gamete formation. Together these results demonstrated that RBPm1 is essential for male gametogenesis and mosquito transmission of parasites.

### Defective axoneme assembly in RBPm1-deficient male gametogenesis

Next we delineated more detailed defects of Δ*Rbpm1*. During male gametogenesis, the parasites undergo axoneme assembly, genome replication, rupture of the parasitophorous vacuole membrane (PVM) and erythrocyte membrane (EM), and finally releasing eight uniflagellated male gametes. We first assessed the axoneme assembly. At 0 mpa, both α- and β-Tubulin were distributed evenly at cytosol of male gametocytes of WT and Δ*Rbpm1* (Figure 3A, upper panel). Immunoblot also detected comparable level for both Tubulins in gametocytes between WT and Δ*Rbpm1* (Figure 3B). At 8 mpa, the axonemal microtubules (MTs) were observed to be coiled around the enlarged nucleus in WT gametocytes. However, aberrant axonemes were formed in Δ*Rbpm1* (Figure A, middle panel). At 15 mpa, Δ*Rbpm1* failed to produce flagellated male gametes (Figure 3A, lower panel). Under ultrastructure expansion microscopy (U-ExM) [34], the axonemes lost bundled structures at 8 mpa in Δ*Rbpm1* compared to the organized axonemes in WT (Figure 3C). We used electron microscope to dissect the ultrastructural defects of axoneme in Δ*Rbpm1* male gametocytes at 8 mpa. The majority of axonemes (93%, 150 axonemes from 43 section images) displayed 9+2 arrangement of MTs in WT (Figure 3D, d1 and d4). In contrast, no intact axoneme (from 69 section images) was detected in either longitudinal or cross sections of Δ*Rbpm1* (Figure 3D, d2 and d5). All the axonemes in Δ*Rbpm1* showed severe defects with loss of either central singlet MTs or peripheral doublet MTs (Figure 3D), consistent with the observation of Tubulin staining in Figure 3C. In the complemented line *rescue*, the axoneme assembly restored normal as in WT (Figure 3D, d3 and d6). These results demonstrated that RBPm1 is required for axoneme assembly during male gametogenesis.

**Figure 3.**
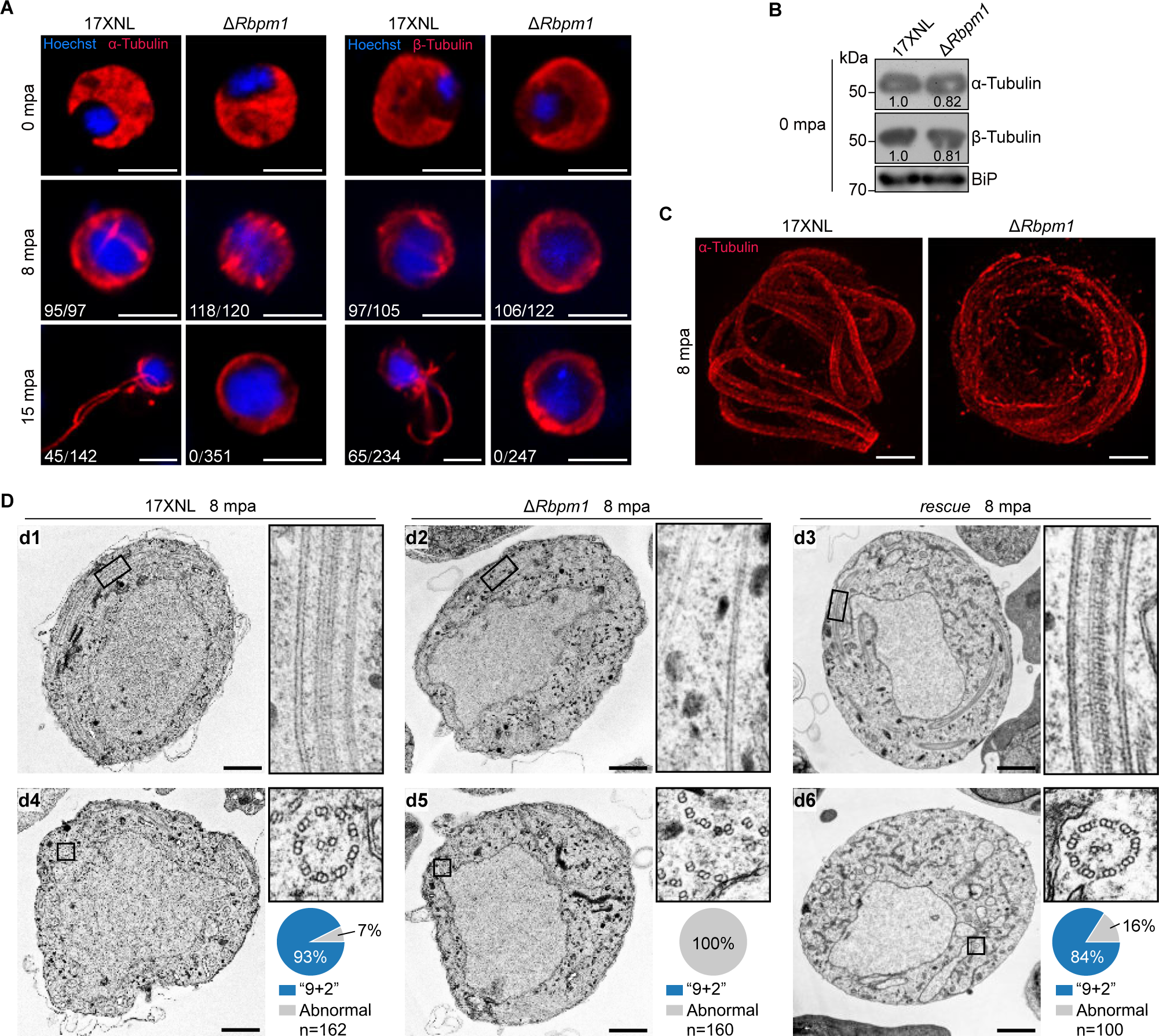
Defective axonemal assembly in RBPm1-null male gametogenesis. **A.** Detection of formation and exflagellation of axonemes during male gametogenesis (0, 8, and 15 mpa) by staining α-Tubulin (left panels) and β-Tubulin (right panels). Nuclei were stained with Hoechst 33342. Representative for three independent experiments. Scale bars: 5 µm. **B.** Immunoblot of α- and β-Tubulins in gametocytes. The numbers indicate the relative intensities of the bands in the immunoblots. BiP as a loading control. Representative for two independent experiments. **C.** Ultrastructure expansion microscopy (U-ExM) of the axonemes in male gametocytes stained with α-Tubulin antibody at 8 mpa. Representative for three independent experiments. Scale bars: 5 μm. **D.** Transmission electron microscopy of axoneme architecture in male gametocytes at 8 mpa. Longitudinal sections (d1, d2, d3 at the top panels) and cross sections (d4, d5, d6 at the bottom panels) of axonemes were shown. The enclosed area (black box) was zoomed in. Pie charts show the quantification of axoneme (“9+2” microtubules) in the mutant parasites. n is the total number of the intact and defective axoneme structures observed in each group. Three independent experiments. Scale bars: 1 μm.

We additionally analyzed genome replication and erythrocyte rupture during male gametogenesis. Flow cytometry analysis of male gametocytes at 8 mpa detected a comparable increase in DNA content in both parental *DFsc7* and its derived mutant *DFsc7*;*ΔRbpm1* parasites (Fig S2A). These results indicated normal genome replication in the absence of RBPm1, consistent with the enlarged nucleus observed in the activated Δ*Rbpm1* male gametocytes from both the fluorescence or electron microscope images (see Figure 3A and D). In addition, protein staining of SEP1 (parasite PVM protein) and TER119 (mouse EM protein) showed that RBPm1 deficiency had no effect on parasite rupture from the gametocyte-residing erythrocytes (Fig S2B and C).

### RBPm1 deficiency causes defective intron splicing of axonemal genes

To investigate the mechanism of RBPm1 in regulating the axoneme assembly, we performed RNA-seq to exam the changes in male transcriptome due to the loss of RBPm1 (Figure 4A). To purify the RBPm1-null male gametocytes for comparison, we deleted *Rbpm1* in the *DFsc7* line. The mutant line *DFsc7*;Δ*Rbpm1* displayed the same phenotypes as Δ*Rbpm1* (Fig S3A-D). Purified male gametocytes of the *DFsc7*;Δ*Rbpm1* were collected by fluorescence-activated cell sorting for RNA-seq (Fig S3E and F). We analyzed the differentially expressed genes between *DFsc7* and *DFsc7*;Δ*Rbpm1*. As expected, the *Rbpm1* transcripts were undetectable in the *DFsc7*;Δ*Rbpm1* (Fig S3G and H). RBPm1 deficiency led to changed expression of many genes, (Fig S3G), but none of the differentially expressed genes was known to be implicated in axoneme assembly during male gametogenesis.

**Figure 4.**
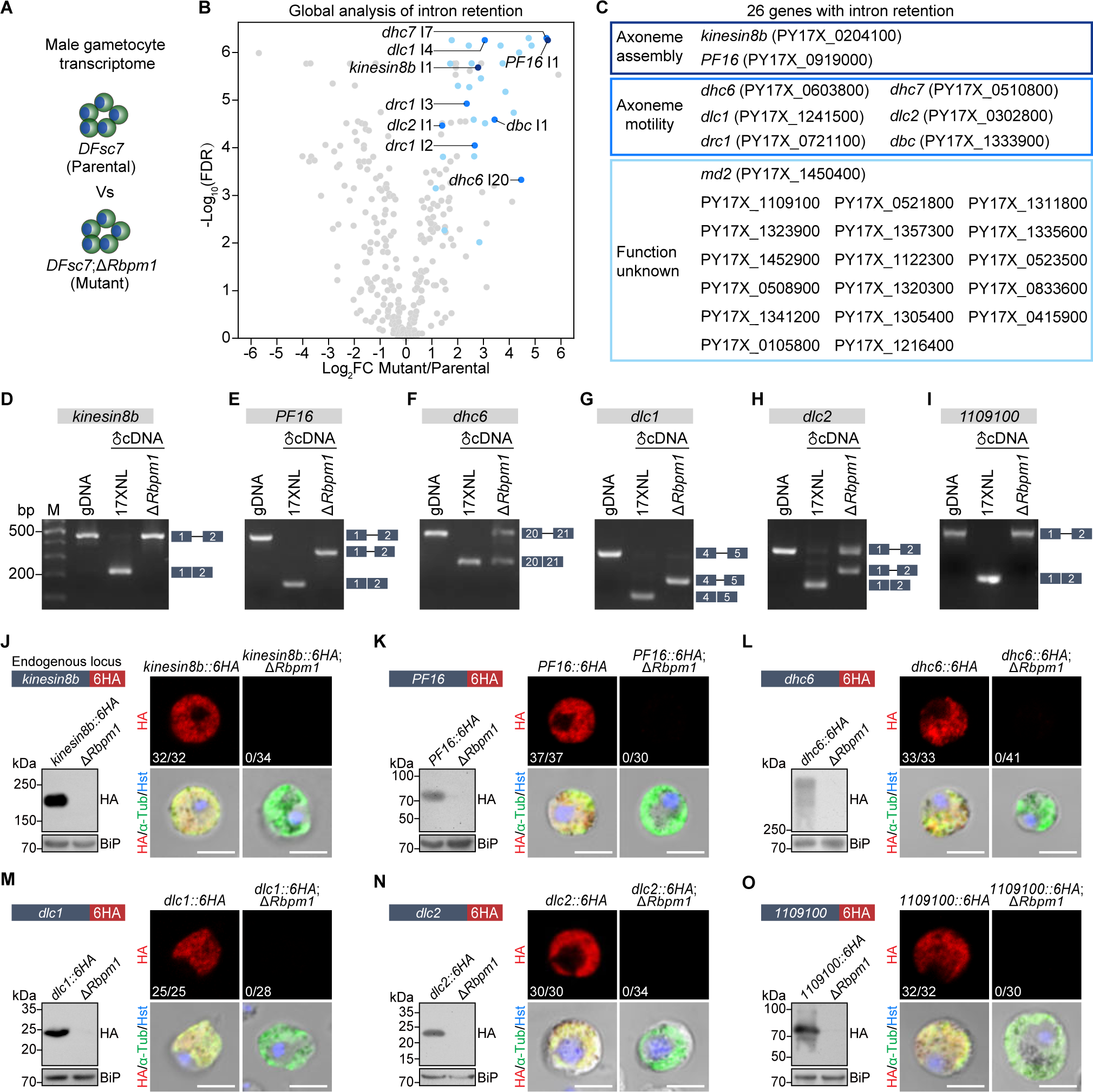
RBPm1 deficiency causes intron retention and protein loss of axonemal genes. **A.** A schematic showing the transcriptome analysis of RBPm1-null male gametocytes. *DFsc7*;Δ*Rbpm1* was a *DFsc7*-derived RBPm1 mutant line. *DFsc7* (parental) and *DFsc7*;Δ*Rbpm1* (mutant) male gametocytes were sorted by FACS for RNA-seq. **B.** Differential intron retention analysis identified 30 retained introns (blue dots) in the mutant versus the parental line. These introns were originated from 26 genes. Detailed information of these genes and introns is provided in Figure S4A. **C.** List of the 26 genes with intron retention from **B**, categorized by protein function. **D-I.** RT-PCR confirmation of intron retention in 6 selected genes *kinesin8b*, *PF16*, *dhc6*, *dlc1*, *dlc2*, and *PY17X_1109100*. Genomic DNA (gDNA) from 17XNL parasite, complementary DNA (cDNA) from male gametocytes of parental and mutant parasites were analyzed. Exons are indicated by boxes and introns by lines. Three independent experiments. RT-PCR analysis for all 26 genes is presented in Figure S5A. **J-O.** Protein expression analysis of the 6 genes shown in **D-I** in male gametocytes after loss of RBPm1. Each endogenous gene was tagged with a 6HA at the C-terminus in both 17XNL and Δ*Rbpm1* parasites (the schematic in the top left panel), generating two tagged lines. Immunoblot of the 6HA-tagged protein in gametocytes with and without RBPm1 (bottom left panel). IFA of the 6HA-tagged protein in male gametocytes with and without RBPm1 (right panel). x/y at bottom-left represents the number of HA-positive male gametocytes/the total number of male gametocytes tested. Representative for three independent experiments. Scale bars: 5 µm.

We found 30 intron retention (IR) events in transcripts of 26 genes after loss of RBPm1 (Figure 4B and C) by bioinformatic analysis of global intron retention and manual examination on Integrative Genomics Viewer [35]. These genes were specifically or preferentially transcribed in male gametocytes compared to female (Fig S4A). Among them (Figure 4C), the orthologs of *kinesin8b* and *PF16* had been reported essential for axoneme assembly in *P. berghei* male gametogenesis [36-38]. Six putative dynein motor-associated genes, *dhc6* (dynein heavy chain, PY17X_0603800), *dhc7* (dynein heavy chain, PY17X_0510800), *dlc1* (dynein light chain, PY17X_1241500), *dlc2* (dynein light chain, PY17X_0302800), *drc1* (dynein regulatory complex protein, PY17X_0721100), and *dbc* (dynein beta chain, PY17X_1333900), were included. The *md2* (male development protein 2, PY17X_1450400), a potential male determining gene recently identified [39], was also included. The rest 17 IR genes had not been previously described in the *Plasmodium*. Gene Ontology (GO) enrichment analysis of these IR genes found significant GO terms that are associated with MT or cytoskeleton (Fig S4B). RT-PCR using the primers anchored in the flank exons of each of 26 introns further confirmed that these introns were retained in the transcripts in the absence of RBPm1, while their neighboring introns were correctly removed (Figure 4D-I and Fig S5A). Using RT-qPCR, we further confirmed the IR of *kinesin8b* intron1 and *PF16* intron1 in the RBPm1-null male gametocytes (Fig S5B). Interestingly, the whole part of intron was retained in the transcripts for most IR genes, while only a N-terminal part of intron was retained for three IR genes, including *PF16* intron1, *dlc1* intron4, and PY17X_1311800 intron5 (Figure 4D-I and Fig S5A). Therefore, RBPm1 is required for the splicing of selective introns in certain male genes, especially MT or cytoskeleton-related genes.

Given the facts that 1. RBPm1 depletion causes defective axoneme assembly, 2. All IR genes are male-specific, 3. Eight IR genes are axoneme-related, we speculated that the RBPm1-regulated IR genes are axonemal. To test it, we selected 12 out of the 26 genes, including 6 annotated (*Kinesin8b*, *PF16*, *dhc6*, *dhc7*, *dlc1*, *dlc2*) and 6 unannotated (PY17X_1109100, PY17X_0521800, PY17X_1311800, PY17X_1323900, PY17X_1357300, PY17X_1335600). Each gene was endogenously tagged at the N- or C-terminus with a 6HA in the 17XNL, generating the HA-tagged lines. All 12 proteins were specifically expressed in male gametocytes during parasite life cycle (Fig S6A), in agree with their transcript profiling. In inactivated gametocytes, these proteins were distributed at cytoplasm, while after activation 11 of 12 proteins displayed axoneme localization in the flagellating male gametes (Fig S6B-M). These results suggested that RBPm1 controls intron splicing for a group of the axonemal genes.

### Intron retention leads to loss of axonemal protein in RBPm1-null male gametocytes

Nucleotide sequence analysis revealed that IR would result in premature translation and thus cause loss of protein expression for the axonemal genes (Fig S7). To analyze the effect of IR on the axonemal proteins after RBPm1 loss, we deleted *Rbpm1* gene in each of two tagged lines *kinesin8B::6HA* and *PF16::6HA*, obtaining two RBPm1-null lines (Figure 4J and K). In the absence of RBPm1, 6HA-tagged Kinesin8B and PF16 were not detected or under detectable level in male gametocytes compared to the parental counterparts in both IFA and immunoblot (Figure 4J and K). To further confirm the protein loss, we analyzed four other IR genes *dhc6*, *dlc1*, *dlc2*, and PY17X_1109100. Endogenous *Rbpm1* gene was deleted in the four tagged lines (*dhc6::6HA*, *dlc1::6HA*, *dlc2::6HA*, and *1109100::6HA*), obtaining another four RBPm1-null lines (Figure 4L-O). These 6HA-tagged proteins lost expression in the RBPm1-null male gametocytes (Figure 4L-O), similarly as Kinesin8B and PF16 did. These results demonstrated that RBPm1 deficiency causes protein expression loss of target axonemal genes.

To confirm the essential roles of *P. yoelii* Kinesin8B and PF16 in axoneme assembly as reported in *P. berghei* [36-38], we disrupted *kinesin8b* and *PF16* genes in the 17XNL, obtaining mutant lines *Δkinesin8b* and *ΔPF16* (Fig S8A). As expected, depletion of *kinesin8b* or *PF16* blocked or severely impaired male gamete formation respectively (Fig S8B). Both mutants produced no midgut oocyst in the infected mosquitoes (Fig S8C). Ultrastructure analysis of male gametocytes at 8 mpa revealed that the *Δkinesin8b* mutant failed to develop “9+2” axoneme with loss of both central and peripheral MTs, while most of the axonemes lost central MTs (shown as “9+0” or “9+1”) in the *ΔPF16* (Fig S8D and E), corresponding with gene disruption phenotypes in *P. berghei* [36-38]. Therefore, depletion of Kinesin8B or PF16 phenocopies RBPm1 deficiency in axoneme assembly.

### Intron deletion restores axonemal proteins and partially rectifies axoneme assembly defects in RBPm1-null gametocytes

Since IR disrupted the axonemal proteins expression, we tested whether enforced genomic deletion of the retained intron could restore protein expression by bypassing intron splicing at the transcripts in RBPm1-null male gametocytes. The endogenous *kinesin8b* intron1 (239 bp) was removed in the *kinesin8b::6HA*;Δ*Rbpm1* parasite by CRISPR-Cas9 (Figure 5A), generating the intron-null mutant *kinesin8b*Δ*intron1* (*kinesin8b*Δ*I1*). Both IFA and immunoblot revealed that the deletion of intron1 restored Kinesin8B::6HA expression to WT level in the RBPm1-null gametocytes (Figure 5B and C). To further confirm the restoration effect, we tested three other retained introns (*PF16* intron1, *dlc1* intron4, and *PY17X_1109100* intron1). Compared to the parental RBPm1-null parasites, the PF16::6HA and 1109100::6HA proteins in male gametocytes were fully restored (Figure 5D-F and 5J-L) while the Dlc1::6HA was partially restored after removal of the corresponding intron (Figure 5G-I). Expression restoration of these axonemal proteins (Kinesin8b, PF16, Dlc1, and PY17X_1109100) via intron deletion strongly confirmed the causative effect of IR on axonemal protein loss in the absence of RBPm1.

**Figure 5.**
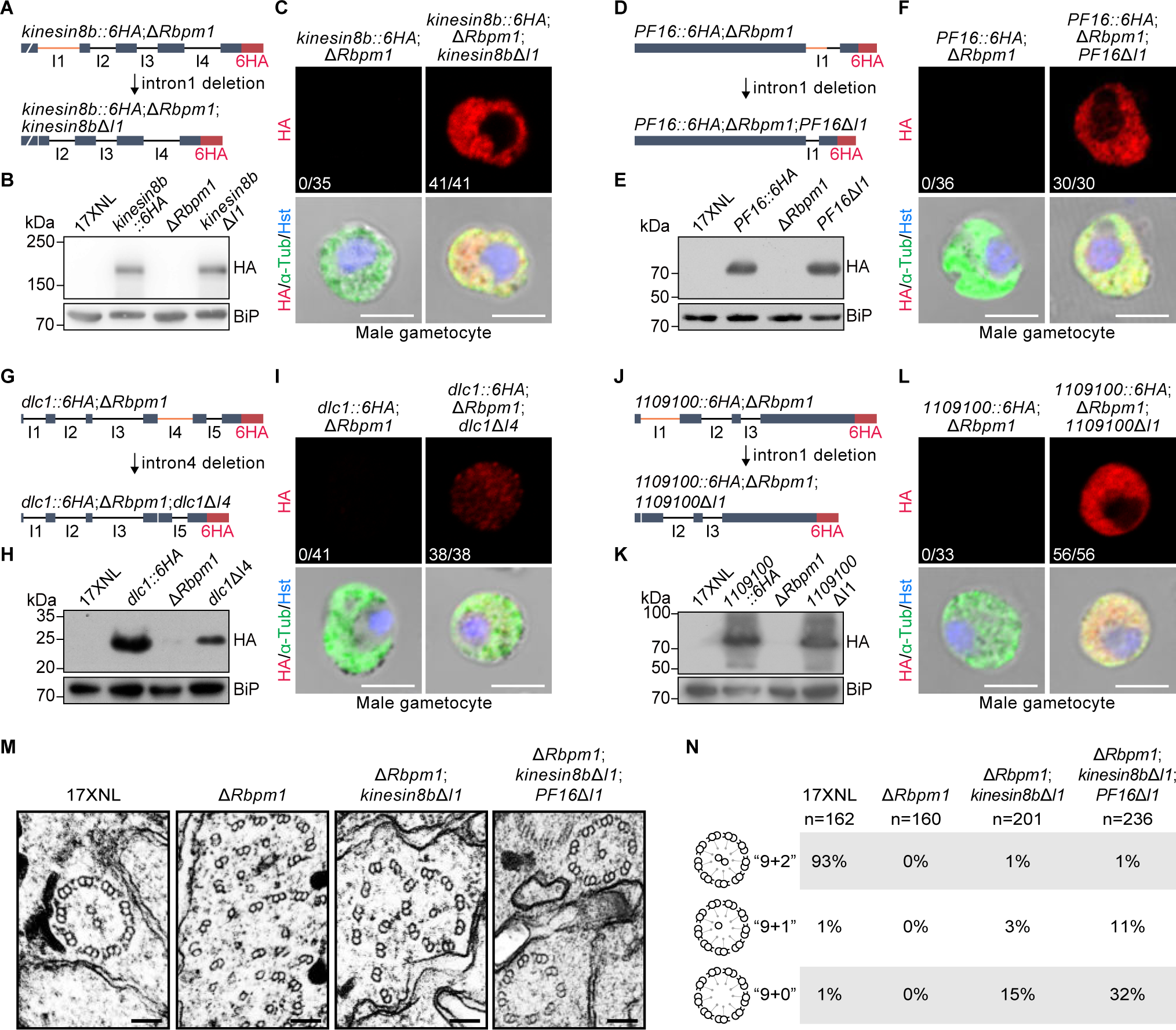
Intron deletion restores axonemal protein expression and partially rectifies axoneme assembly defects in RBPm1-null male gametocytes. **A.** A schematic showing the genomic deletion of retained intron (*kinesin8b* intron1, orange) in the *kinesin8b::6HA*;Δ*Rbpm1* parasite (abbreviated as Δ*Rbpm1*), generating the mutant *kinesin8b::6HA*;Δ*Rbpm1*;*kinesin8b*Δ*intron1* (abbreviated as *kinesin8b*Δ*I1*). **B.** Immunoblot of 6HA-tagged Kinesin8B protein in gametocytes. Representative for three independent experiments. **C.** IFA of 6HA-tagged Kinesin8B in male gametocytes. x/y represents the number of HA-positive male gametocytes/the total number of male gametocytes tested. Representative for three independent experiments. Scale bars: 5 µm. **D, E, and F.** Effect of intron deletion (*PF16* intron1) on the restoration of PF16 protein in RBPm1-null male gametocytes. Similar analysis as in **A**. **B**. and **C**. **G, H, and I.** Effect of intron deletion (*dlc1* intron4) on the restoration of Dlc1 protein in RBPm1-null male gametocytes. **J, K, and L.** Effect of intron deletion (*PY17X_1109100* intron1) on the restoration of PY17X_1109100 protein in RBPm1-null male gametocytes. **A. M.** Transmission electron microscopy of axoneme architecture in male gametocytes at 8 mpa. Δ*Rbpm1*;*kinesin8b*ΔI1 is a Δ*Rbpm1*-derived modified line with deletion of *kinesin8b* intron1. Δ*Rbpm1*;*kinesin8b*ΔI1;*PF16*ΔI1 is a Δ*Rbpm1* derived modified line with deletion of both *kinesin8b* intron1 and *PF16* intron1. Scale bars: 100 nm. **B. N.** Quantification of axoneme formation from parasites in **M**. n is the total number of the intact and defective axoneme structures observed in each group. Representative for three independent experiments.

We next tested whether genomic deletion of the retained introns could rescue or rectify the defective axoneme assembly in the Δ*Rbpm1* mutant. We deleted the *kinesin8b* intron1 in the Δ*Rbpm1* line, but this deletion of single intron failed to restore any EC formation in the Δ*Rbpm1*;*kinesin8b*Δ*intron1* parasites. However, compared to complete lack of axonemes showing “9+2”, “9+1”, or “9+0” MTs in the parental Δ*Rbpm1*, some axoneme-like structures (“9+2”: 1%, “9+1”: 3%, and “9+0”: 15%) were detected in the Δ*Rbpm1*;*kinesin8b*Δ*intron1* (Figure 5M and N), indicating that deletion of *kinesin8b* intron1 could partially rectify the defective axoneme assembly caused by RBPm1 deficiency. Notably, additional deletion of the *PF16* intron1 in the Δ*Rbpm1*;*kinesin8b*Δ*intron1* parasite further mitigated axoneme defects in the resulted Δ*Rbpm1*;*kinesin8b*Δintron1;*PF16*Δintron1 parasite (“9+2”: 1%, “9+1”: 11%, and “9+0”: 32%) (Figure 5M and N). These results demonstrated that RBPm1 regulates axoneme assembly by controlling intron splicing of a group of axonemal genes. Without RBPm1, deletion of two introns (*kinesin8b* intron1 and *PF16* intron1) was insufficient to restore axoneme assembly to the WT level (“9+2”: 93%) (Figure 5M and N). Therefore, in addition to *kinesin8b* and *PF16*, other axonemal genes targeted by RBPm1 may also play important roles in axoneme assembly during male gametogenesis.

### RBPm1 interacts with spliceosome E complex and introns of axonemal genes

To investigate whether RBPm1 associates with the spliceosome responsible for intron splicing, we used the biotin ligase TurboID-based proximity labeling to identify RBPm1-interacting proteins in the gametocytes. The endogenous RBPm1 was tagged with a HA::TurboID motif in the 17XNL, generating the line *Rbpm1::TurboID* (Figure 6A). A control parasite *Rbpm1::T2A::TurboID* was generated by fusing endogenous RBPm1 with a “ribosome skip” T2A peptide, a NLS (nuclear localization signal), and a HA::TurboID (Figure 6A), permitting separated expression of RBPm1 and biotin ligase. Gametocytes expressing the ligase were incubated with 50 µM biotin for 20 min at 37°C. Staining with fluorescent-conjugated streptavidin and anti-HA antibody detected a nuclear distribution of biotinylated proteins in both TurboID-modified gametocytes (Fig S9A), indicating biotinylation of the potential RBPm1-interacting proteins in the nucleus. Mass spectrometry of the streptavidin affinity purified proteins from the *Rbpm1::TurboID* resulted in a list of 113 proteins enriched with high confidence compared to the control (Figure 6B). RBPm1 was the top hit, confirming cis-biotinylation of RBPm1 (Figure 6B). Among the significantly enriched proteins, we found the components of the spliceosome earliest assembling E complex [40-42], including the U1 small nuclear ribonucleoproteins (snRNP) U1-70K, U1-A, U1-C, Sm-B, Sm-D1, Sm-D2, Sm-D3, Sm-E, Sm-F and Sm-G (Figure 6B and C), and three E complex key factors SF1, U2AF1, and U2AF2 (Figure 6B and C). Tagging the endogenous U1-70K, U1-A and U1-C proteins with 4Myc in the *Rbpm1::6HA* parasite showed that these three U1 snRNPs co-localized with RBPm1 in the nucleus (Figure 6D). Co-immunoprecipitation also confirmed the interaction between RBPm1 and these U1 snRNPs (Figure 6E-G). Spliceosome A, B, and C complex are formed after the assembly of splicing initiating E complex [43, 44]. However, the components of A, B, and C complex were not detected (Fig S9B and C). Therefore, RBPm1 interacted only with spliceosome E complex, possibly helping to initiate splicing for certain introns in the axonemal genes (Figure 6H).

**Figure 6.**
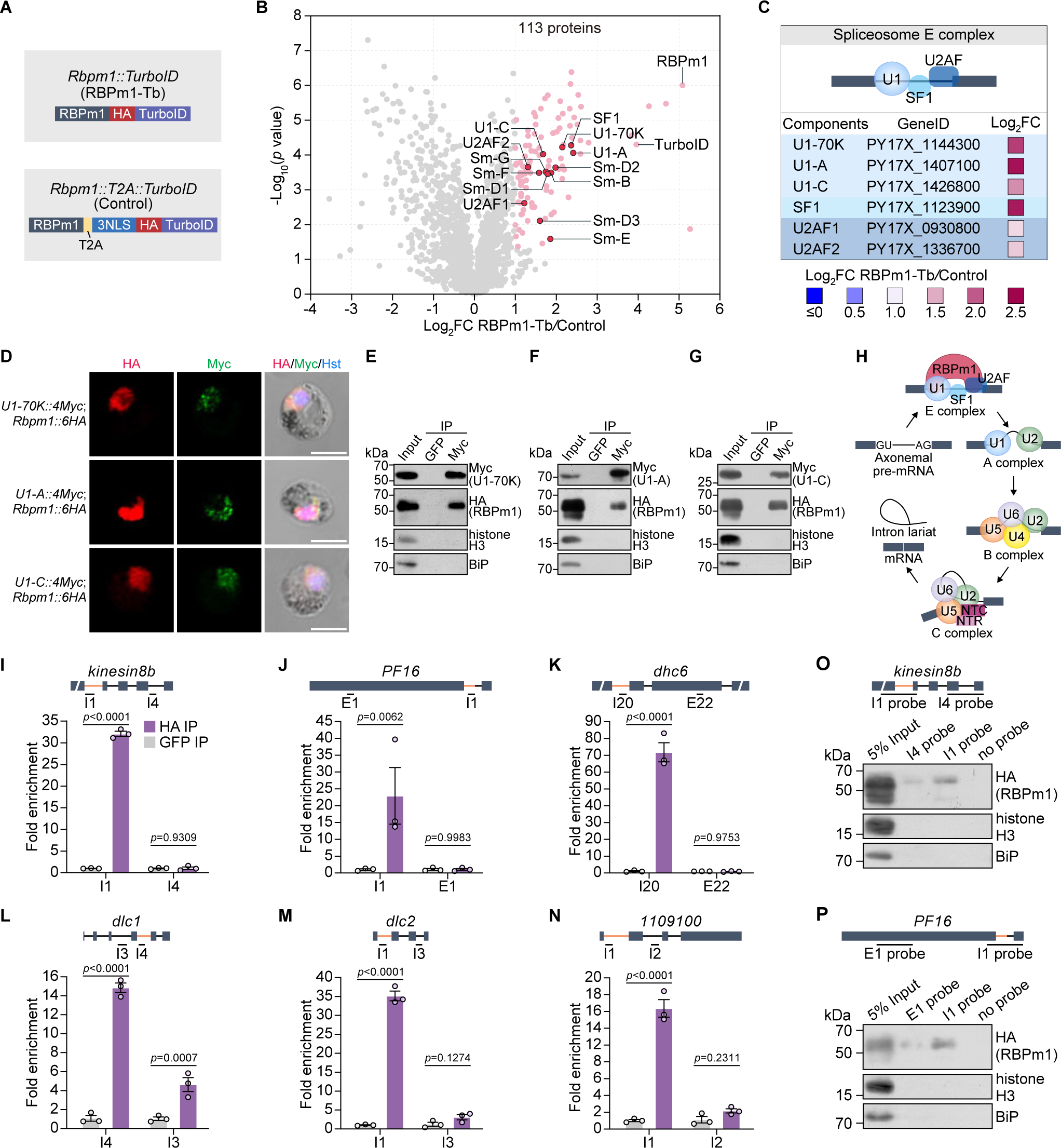
RBPm1 interacts with spliceosome E complex and introns of axonemal genes. **A.** A schematic showing two modified parasite lines generated for searching RBPm1-interacting proteins by TurboID-based proximity labeling and mass spectrometry. The motif of HA::TurboID and T2A::3NLS::HA::TurboID, respectively, was inserted at the C-terminus of the endogenous RBPm1, generating the line *Rbpm1::TurboID* and the control line *Rbpm1::T2A::TurboID*. **B.** Volcano plot displaying 113 significantly enriched proteins (pink dot, cutoffs log_2_FC ≥ 1 and *p*-value ≤ 0.05) in the *Rbpm1::TurboID* versus *Rbpm1::T2A::TurboID*. Among them, 13 subunits (red dot) of the early spliceosome E complex were included. **C.** Protein interaction analysis between RBPm1 and six spliceosome E complex subunit proteins (U1-70K, U1-A, U1-C, SF1, U2AF1, and U2AF2) from **B**. **D.** IFA of 6HA-tagged RBPm1 and 4Myc-tagged U1 snRNP proteins (U1-70K, U1-A, and U1-C) in male gametocytes of three double-tagged parasites. Representative for three independent experiments. Scale bars: 5 µm. **E.** Co-immunoprecipitation of RBPm1 and U1-70K in gametocytes of the double-tagged parasite *U1-70K::4Myc*;*Rbpm1::6HA*. Anti-Myc nanobody was used. Bip as a loading control. Representative for three independent experiments. **F.** Co-immunoprecipitation of RBPm1 and U1-A in gametocytes of the double-tagged parasite *U1-A::4Myc*;*Rbpm1::6HA*. Three independent experiments. **G.** Co-immunoprecipitation of RBPm1 and U1-C in gametocytes of the double-tagged parasite *U1-C::4Myc*;*Rbpm1::6HA*. Three independent experiments. **H.** Proposed model showing the interaction between RBPm1 and early spliceosome E complex for intron splicing of axonemal genes. **I-N.** UV-RIP detection of RBPm1 interaction with the retained introns of 6 axonemal genes (*kinesin8b*, *PF16*, *dhc6*, *dlc1*, *dlc2*, and *PY17X_1109100*). A top schematic shows the exon-intron structure of the RBPm1 target axonemal genes. The retained introns are indicated with orange lines, and the genomic regions for qPCR amplicon are shown. UV-RIP was performed in *Rbpm1::6HA* lines using anti-HA nanobody. Anti-GFP nanobody was used as a control. Bound RNA was analyzed by RT-qPCR. Means ± SEM from three independent experiments, two-tailed *t*-test. **O.** RNA pull-down assay detecting RBPm1 interaction with *kinesin8b* intron1. A top schematic shows the exon-intron structure of the *kinesin8b* gene. The retained introns are indicated in orange lines. A biotinylated 500 nt RNA probe I1 (comprising intron1 and its flanking sequences) and a control probe I4 (comprising intron4 and its flanking sequences) were used. Proteins via RNA pull-down were immunoblot with anti-HA antibody. Histone H3 and Bip were used as negative controls. Two independent experiments with similar results. **P.** RNA pull-down assay detecting RBPm1 interaction with *PF16* intron1. A top schematic shows the exon-intron structure of the *PF16* gene. The retained introns are indicated in orange lines. A biotinylated 500 nt RNA probe I1 (comprising intron1 and its flanking sequences) and a control probe E1 in exon1 were used. Two independent experiments with similar results.

Nuclear localization and interaction with spliceosome E-complex imply that RBPm1 may bind to the target introns in the pre-mRNA of axonemal genes. We performed UV crosslinking RNA immunoprecipitation (UV-RIP) followed by RT-qPCR with primers recognizing the target pre-mRNAs. In the *Rbpm1*::*6HA* gametocytes, RBPm1 bound to the intron1 of the *kinesin8b* transcripts using anti-HA nanobody (Figure 6I). As a control, RIP using anti-GFP nanobody detected no binding (Figure 6I). Additionally, we analyzed the interaction between RBPm1 and five other target introns (*PF16* intron1, *dhc6* intron20, *dlc1* intron4, *dlc2* intron1, and *PY17X_1109100* intron1). As expected, RBPm1 bound these target introns but not the neighboring introns or exons (Figure 6J to N).

Furthermore, we used RNA pull-down to validate the interaction of RBPm1 with the *kinesin8b* intron1 and *PF16* intron1. A biotinylated 500nt RNA probe *kinesin8b* I1 and a control probe *kinesin8b* I4 were synthesized (Figure 6O, upper panel) and incubated with the *Rbpm1::6HA* gametocyte lysate. The potential RNA-interacted proteins were precipitated using the streptavidin beads and detected by immunoblot. The *kinesin8b* I1 probe retrieved more RBPm1 protein than the *kinesin8b* I4 probe (Figure 6O). Similarly, the *PF16* I1 probe captured more RBPm1 protein than the probe *PF16* E1 (Figure 6P). Both RIP and RNA pull-down experiments supported that RBPm1 binds the *kinesin8b* intron1 and *PF16* intron1.

### RBPm1 directs splicing of axonemal introns inserted in a reporter gene

To further investigate the interaction between RBPm1 and the axonemal introns, we test whether RBPm1 could direct splicing of target introns when inserted into a reporter gene. We developed a blue fluorescence protein (BFP) reporter assay that allows an easy splicing readout in male and female gametocytes of the *DFsc7* parasite. The intact *bfp* transcript driven by the *hsp70* 5’-UTR and the *dhfr* 3’-UTR was integrated into the *p230p* locus of *DFsc7* using CRISPR-Cas9, generating the control line *BFP* (Figure 7A). The *kinesin8b* intron1 (*Kin8b*I1, 239 bp) was inserted to the *bfp* gene at the nucleotides 396-397, generating the line *BFP*-*Kin8b*I1 (Figure 7B). The inserted *kinesin8b* intron1 would result in premature translation of the *bfp* transcript if it is not spliced. In the control *BFP* line, BFP was expectedly detected in both male (GFP+) and female (mCherry+) gametocytes (Figure 7A). However, in the *BFP*-*Kin8b*I1 line, BFP was detected only in male gametocytes (Figure 7B), indicating that splicing of *kinesin8b* intron1 in the *bfp* transcripts occurred only in male gametocytes. To prove that splicing of *kinesin8b* intron1 in male was RBPm1-dependent, we deleted *Rbpm1* in the *BFP*-*Kin8b*I1 line and obtained the mutant line *BFP*-*Kin8b*I1;Δ*Rbpm1* (Figure 7C). RBPm1 deletion disrupted BFP expression in the *BFP*-*Kin8b*I1;Δ*Rbpm1* male gametocytes (Figure 7C). We parallelly analyzed the *kinesin8b* intron2 (*Kin8b*I2, 148 bp), whose splicing from the native gene transcript required no RBPm1 (Fig S5). In both transgenic line *BFP*-*Kin8b*I2 (Figure 7D) and its derived mutant line *BFP*-*Kin8b*I2;Δ*Rbpm1* (Figure 7E), BFP was detected in both male and female gametocytes, confirming RBPm1-independent splicing of *kinesin8b* intron2 from the *bfp* transcript.

**Figure 7.**
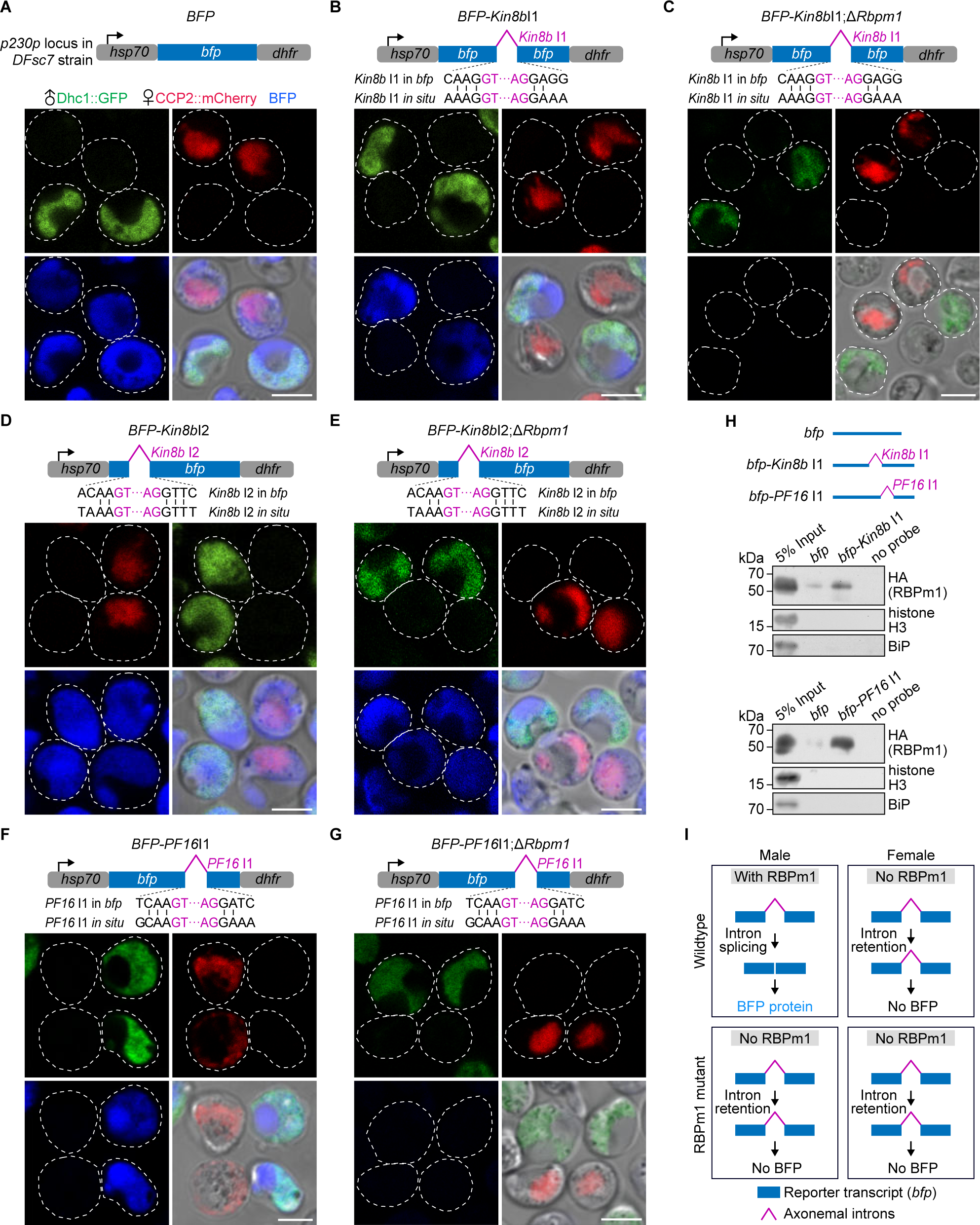
RBPm1 directs splicing of axonemal introns inserted in a reporter gene. **A.** A top schematic shows a transgenic line *BFP* with a *bfp* reporter expression cassette integrated at the *p230p* locus of the *DFsc7* reporter line. The intact *bfp* is driven by the 5′UTR of *hsp70* and the 3′UTR of *dhfr*, allowing expression of BFP in both male (GFP+) and female (mCherry+) gametocytes. Live cell imaging was shown. Representative for three independent experiments. Scale bars: 5 µm. **B.** A transgenic line *BFP-Kin8b*I1 with a *kinesin8b* intron 1 (*Kin8b* I1, purple line)-inserted *bfp* cassette integrated at the *p230p* locus of the *DFsc7* line. *Kin8b* I1 (purple) was inserted to the *bfp* gene at the nucleotides 396-397 to mimic the splice site (vertical lines) of *in situ Kin8b* I1. BFP expression was detected specifically in male gametocytes of the *BFP-Kin8b*I1 parasites. Three independent experiments. Scale bars: 5 µm. **C.** A *BFP-Kin8b*I1 derived RBPm1 mutant line *BFP-Kin8b*I1;Δ*Rbpm1*. No BFP expression was detected in male gametocytes of the *BFP-Kin8b*I1;Δ*Rbpm1* parasites. Three independent experiments. Scale bars: 5 µm. **D.** Effect of the *kinesin8b* intron 2 (*Kin8b* I2) insertion on the gametocyte expression of BFP. Similar analysis as in **B**. BFP expression was detected in both male and female gametocytes of the *BFP-Kin8b*I2 parasites. **E.** A *BFP-Kin8b*I2 derived RBPm1 mutant line *BFP-Kin8b*I2;Δ*Rbpm1*. Similar analysis as in **C**. BFP expression was detected in both male and female gametocytes of the *BFP-Kin8b*I2;Δ*Rbpm1* parasites. **F.** Effect of the *PF16* intron1 (*PF16* I1) insertion on the gametocyte expression of BFP. Similar analysis as in **B**. BFP expression was detected specifically in male gametocytes of the *BFP-PF16*I1 parasites. **G.** A *BFP-PF16*I1 derived RBPm1 mutant line *BFP-PF16*I1;Δ*Rbpm1*. Similar analysis as in **C**. No BFP expression was detected in male gametocytes of the *BFP-PF16*I1;Δ*Rbpm1* parasites. **H.** RNA pull-down assay detecting RBPm1 interaction with the *Kin8b* I1 and *PF16* I1 -inserted *bfp* transcripts from *BFP-Kin8b*I1 and *BFP-PF16*I1 gametocytes, respectively. Three biotinylated RNA probes *bfp*, *bfp-Kin8b*I1 (corresponding to the *Kin8b* I1-inserted *bfp* transcript), *bfp-PF16*I1 (corresponding to the *PF16* I1-inserted *bfp* transcript) were used. Proteins via RNA pull-down were immunoblotted with anti-HA antibody. Histone H3 and Bip were used as negative controls. Two independent experiments with similar results. **I.** A schematic of RBPm1-dependent splicing of axonemal introns inserted in the reporter transcript.

Using the reporter assay, we tested 3 other target introns, including *PF16* intron1 (Figure 7F and G), *dlc1* intron4 (Fig S10A and B), and *PY17X_1109100* intron1 (Fig S10C and D). The *PF16* intron1 (276 bp) was inserted to the *bfp* at the nucleotides 500-501 (Figure 7F), the *dlc1* intron4 (193 bp) at the nucleotides 455-456 (Fig S10A), while the *PY17X_1109100* intron1 (353 bp) at the nucleotides 390-391 (Fig S10C). As expected, these introns were spliced from the *bfp* transcript only in male gametocytes (Figure 7F, Fig S10A and C). Similarly, these introns were not spliced at male gametocytes in the RBPm1-null parasites compared to their parental parasites (Figure 7G, Fig S10B and D). Additionally, we analyzed the *PY17X_1109100* intron2 (272 bp), which could be spliced in the RBPm1-null parasites (Fig S5), and found that RBPm1 was not required for splicing of this intron from *bfp* transcript in both male and female gametocytes (Fig S10E and F).

Furthermore, we analyzed the RBPm1 interaction with the *kinesin8b* intron1 and *PF16* intron1 in the *bfp* transcript by RNA pull-down. A biotinylated RNA probe *bfp-Kin8b*I1, corresponding to the *kinesin8b* intron1-inserted *bfp* transcript (Figure 7H, upper panel), retrieved significantly more RBPm1 from the *Rbpm1::6HA* gametocyte lysate compared to the control probe *bfp* (Figure 7H, middle panel). Similarly, the probe *bfp-PF16*I1, corresponding to the *PF16* intron1-inserted *bfp* transcript, captured more RBPm1 than the control probe *bfp* (Figure 7H, lower panel). Therefore, RBPm1 could recognize the axonemal introns in the reporter transcript for splicing (Figure 7I).

### RBPm1 directs splicing of axonemal introns inserted in an endogenous gene

In addition to the reporter gene, we also tested whether RBPm1 could direct splicing of target introns when inserted into an endogenous gene which does not require RBPm1 for intron splicing. We chose the *gep1*, a 4-exon gene expressed in both gender gametocytes and essential for initiating both genders’ gametogenesis [45]. We analyzed male and female gametogenesis by measuring EM rupture (TER119 staining), genome replication (DNA staining), and axoneme assembly (α-Tubulin staining). Compared to 17XNL, the *gep1*-deleted parasite line *Δgep1* expectedly lost ability in EM rupture, genome replication, and axoneme assembly in activated male gametocytes, as well as EM rupture in activated female gametocytes (Figure 8A, B, E, F, G, and H). Using CRISPR-Cas9, the *kinesin8b* intron1 was inserted into the exon3 of *gep1* locus at the nucleotides 273-274 in the 17XNL (Figure 8C), while the *PF16* intron1 inserted into the exon1 at the nucleotides 885-886 (Figure 8D). In both intron-inserted lines *gep1*-*Kin8b*I1 and *gep1*-*PF16*I1, normal male gametogenesis and defective female gametogenesis was speculated because GEP1 is not expressed in female due to no RBPm1-mediated intron splicing from the *gep1* transcript. Notably, both the *gep1*-*Kin8b* I1 and *gep1*-*PF16* I1 parasites underwent EM rupture only in male (Figure 8C-F). These results supported that both *kinesin8b* intron1 and *PF16* intron1 were spliced from the *gep1* transcript only in male gametocytes with RBPm1 expression. Consistent with GEP1 expression in male gametocytes, normal genome replication and axoneme assembly were detected during male gametogenesis in both *gep1*-*Kin8b* I1 and *gep1*-*PF16* I1 parasites (Figure 8C, D, G and H). Collectively the results from the reporter and the endogenous gene assays (Figure 8I) indicated that the tested axonemal introns themselves could be specifically recognized by RBPm1 for splicing.

**Figure 8.**
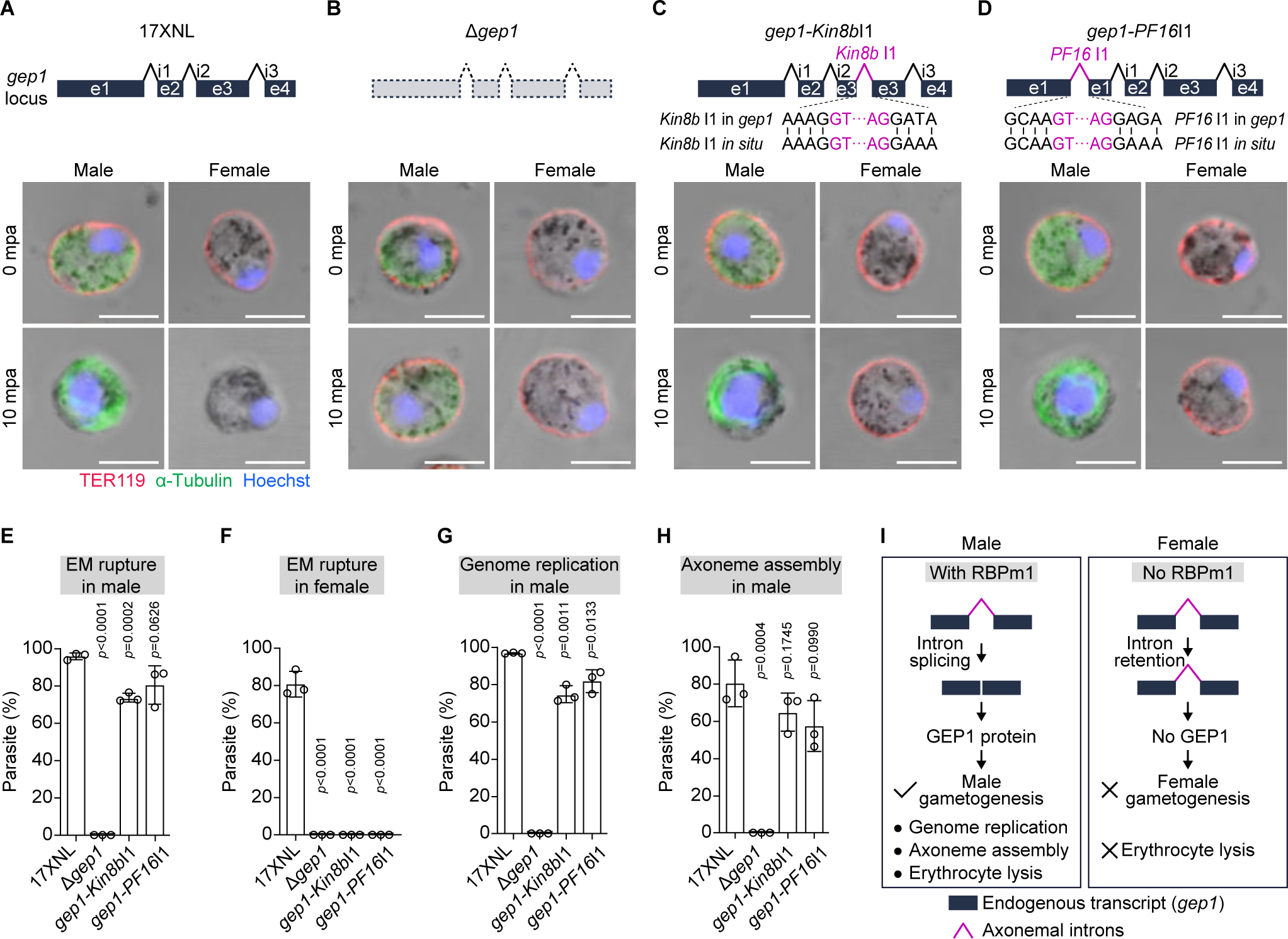
RBPm1 directs splicing of axonemal introns inserted in the endogenous gene. **A.** A top schematic shows the genomic locus of a 4-exon gene *gep1*, which is expressed in both gender gametocytes and essential for both genders’ gametogenesis. Erythrocyte plasma membrane (EM) rupture, genome replication, and cytoplasmic assembly of the axoneme were analyzed in gametocytes of the 17XNL parasites at 10 mpa. The parasites were co-stained with anti-TER-119 and anti-α-Tubulin antibodies and Hoechst 33342. TER-119 (red) negative gametocytes were recognized as EM rupture. Enlarged nuclei represent the genome replication in male gametocytes. Enhanced α-Tubulin (green) signal represents cytoplasmic assembly of the axoneme in male gametocytes. Representative for three independent experiments. Scale bars: 5 µm. **B.** A top schematic shows a modified line Δ*gep1*, in which the endogenous *gep1* gene was deleted in the 17XNL. Similar analysis for the Δ*gep1* parasites as in **A**. **C.** A top schematic shows a modified line *gep1*-*Kin8b*I1, in which the *kinesin8b* intron1 (*Kin8b*I1) was inserted into the exon3 of *gep1* locus at the nucleotides 273-274 in the 17XNL. Similar analysis for the *gep1*-*Kin8b*I1 parasites as in **A**. **D.** A top schematic shows a modified line *gep1*-*PF16*I1, in which the *PF16* intron1 (*PF16*I1) was inserted into the exon1 of *gep1* locus at the nucleotides 885-886 in the 17XNL. Similar analysis for the *gep1*-*PF16*I1 parasites as in **A**. **E.** Quantification of EM rupture in male gametocytes of the four parasites tested. Data are means ± SEM from three independent experiments, two-tailed *t*-test. **F.** Quantification of EM rupture in female gametocytes of the four parasites tested. Data are means ± SEM from three independent experiments, two-tailed *t*-test. **G.** Quantification of genome replication in male gametocytes of the four parasites tested. Data are means ± SEM from three independent experiments, two-tailed *t*-test. **H.** Quantification of axoneme assembly in male gametocytes of the four parasites tested. Data are means ± SEM from three independent experiments, two-tailed *t*-test. **I.** Schematic of RBPm1-dependent splicing of axonemal introns inserted in the endogenous gene *gep1*. Male-specific RBPm1 could recognize and splice the axonemal introns (*kinesin8b* intron1 or *PF16* intron1) inserted in the *gep1* transcript, allowing male-specific GEP expression and thus male gametogenesis.

## Discussion

For efficient transmission, the malaria parasites in the vertebrate host differentiate into sexual precursor male and female gametocytes that are poised to rapidly activate to fertile gametes upon entering into the midgut for fertilization and further development. So far, a limited number of transcription and epigenetic factors have been identified during gametocyte and gamete development, suggesting the post-transcriptional regulation in these transition processes [46, 47]. Consistently, among approximately 180 putative *Plasmodium* RBPs, about one-third of RBP genes exhibit stage specific or elevated expression at the gametocyte [30]. From this list of RBPs, we identified a previously undescribed male-specific nuclear RBP, RBPm1, which was essential for male gametogenesis and mosquito transmission of the *Plasmodium*. RBPm1 operated as a stage- and gender-specific splicing factor for spliceosome assembly initiation and regulated the protein expression for a group of 26 male genes, most of which are axoneme-related. Recent studies had discovered several other RBPs playing crucial roles in the developmental programs of gametocyte and gamete [48-50]. During gametocytes development, UIS12 contributes to the development of both gender gametocytes [51]. Disrupting *Puf1* led to a reduction in gametocytes, especially female gametocytes [52], while *ccr4-1* gene deletion obstructs male gametocyte development [53]. *Puf2* knockout, on the other hand, promotes male gametocyte development [54]. In female gametocytes, the DOZI/CITH/ALBA translation repressor complex and PUF2 protect the stored mRNAs for translation repression until needed during the development of post fertilization [26, 55]. Additionally, the CAF1/CCR4/NOT complex also plays a role in safeguarding the stored mRNAs from degradation. In male gametocytes, two RBPs have been identified functionally. The alternative splicing factor SR-MG promotes the establishment of sex-specific splicing patterns and knocking it out reduces the male gamete formation [48]; the *ZNF4*’s knockout results in deregulation of 473 genes, including axonemal dynein related genes in mature gametocytes [56]. These documented RBPs and RBPm1 identified in this study may function together at the post-transcription regulation to shape the male distinct transcriptome for male gametocyte and gamete development.

During the manuscript preparation of this study, another work by Russell *et al*. [39] demonstrated that gene deletion of the *Rbpm1* ortholog (PBANKA_0716500, named as *md5*) had no effect on female and male gametocyte formation, but resulted in male-specific infertility in the *P. berghei*. These findings are consistent with the male gamete formation deficiency phenotype of the *ΔRbpm1* in *P. yoelii* in this study, indicating conserved function of RBPm1 in the rodent malaria parasites. As *P. falciparum* is the most lethal human malaria parasite, future studies are worthy to investigate whether the RBPm1 ortholog in the *P. falciparum* functions similarly in male gametogenesis.

The RBPm1-deficient parasites showed specific defects in axoneme assembly during male gametogenesis (Figure 3). Axoneme is a MT cytoskeleton essential for the eukaryotic flagellar motility, consisting of a central pair of singlet MTs encircled by 9 outer doublet MTs. This 9+2 organization of axonemes is highly conserved in the eukaryotes, including *Plasmodium* [57]. However, the axoneme in *Plasmodium* differs from that in model organisms in several aspects [10, 58, 59]. First, the biogenesis of axoneme in *Plasmodium* male gametogenesis is extremely fast, taking only 6-8 minutes to assemble 8 axonemes [9, 10]. Second, location of basal body. In the canonical cilium, the basal body is localized under the plasma membrane. In *Plasmodium* male gametocytes, the basal bodies are residing at the nuclear membrane [59]. Third, location for axoneme assembly. The canonical axoneme protrudes distally from the cell simultaneously when growing from the basal body. The *Plasmodium* assemblies the axoneme completely within the cytoplasm, independent of intraflagellar transport required in the cilium formation [10, 58]. Last, each assembled axoneme associates with a haploid nuclei to progressively protrude from the parasite plasma membrane, resulting in a free motile flagellum [10]. Mechanisms underlying the cytoplasmic assembly and exflagellation of axonemes in *Plasmodium* remain largely unknown, although the involvement of some conserved basal body and axonemal proteins has been described, including the armadillo repeat protein PF16 [38], the motor protein Kinesin8B [36, 37, 60], the basal body proteins SAS4 and SAS6 [59, 61-63], and the radial spoke protein RSP9 [64]. It is reasonable that the *Plasmodium* had evolved novel mechanisms to fulfill the requirement for the axoneme. In this study, we identified a group of 26 male genes targeted by RBPm1. Several known or putative axoneme-associated genes were included. Importantly, endogenous protein localization analysis showed that most of the tested RBPm1-target genes encode proteins co-localizing with axoneme (Fig S6B-M), suggesting their roles in biogenesis, structure, regulation, or function of axoneme. For future studies, it is intriguing to understand the roles of these 26 RBPm1-regulated genes, especially 17 previously undescribed one, during male gametogenesis in the *Plasmodium*.

In the RBPm1 deficient male gametocytes, 30 IR events were detected in 26 male genes. One intron was retained in each of 22 genes while two introns were retained in each of 4 other genes (Figure 4B and Figure S4A). Mechanically, RBPm1 not only bound to the intron-retained transcripts, but also interacted with spliceosome E complex. U1 snRNPs recognize and pair with the 5′ splice site of intron. SF1, U2AF1 and U2AF2 form a complex and bind to branch point, 3′ splice site, and polypyrimidine tract respectively [65, 66]. These above factors assemble the E complex as a spliceosome earliest stage. After that, the spliceosome dynamically releases and recruits different snRNPs to establish the assembly for further stages, including spliceosome A, B, and C complex. RBPm1 was detected to interact exclusively with the components of spliceosome E complex, but not of the A, B, and C complex (Fig S9B and C). Therefore, RBPm1 likely function as a splicing activator, linking spliceosome E complex with the selective introns of axonemal genes for splice site recognition. In mammals and plants, an RBP of Dek played a similar role and promoted the splicing of certain introns by bridging the intron with the U1/U2 snRNPs [67, 68]. At this stage, the data support an association of RBPm1 and spliceosome E complex, but it is not yet clear if it is a direct or indirect association.

Both RIP and RNA pull-down assays demonstrated that RBPm1 interacts with the target introns in transcripts of the axonemal genes, suggesting the presence of signals recognized by RBPm1 in these introns. To investigate that the signals for RBPm1 recognition are afforded by the introns themselves and not by the adjoining exons, we analyzed the splicing capability of these introns when they were inserted in either a reporter gene (*bfp*) or an irrelevant endogenous gene (*gep1*). The results from both intron splicing assays (Figure 7 and 8) exhibit stringent dependencies of splicing on RBPm1 for these axonemal introns, suggesting intrinsic signals within the introns for RBPm1 recognition. In addition, the adjoining exons may play less role in the RBPm1 recognition of the axonemal introns. We attempted to search the common characteristics, including the length, GC content, splice sites, and motif enrichment, but unfortunately observed no distinct features among these 30 RBPm1 target introns. The molecular basis for these axonemal intron recognition by RBPm1 is still unknown. One possibility is that RBPm1 target introns may possess the sequence-independent features, such as RNA structures or epigenetic modifications, for RBPm1 recognition. The structure of RBPm1 is not available yet. To understand the recognition or interaction between RBPm1 and its target introns, future studies into an atomic resolution structure of the protein (RBPm1)-RNA (intron) complex will be required.

Axoneme is an essential cellular structure specifically required in male gametogenesis during the *Plasmodium* life cycle. Consistently, the axonemal genes display significant male-biased transcription [72-74]. From previous and this studies (Fig S11A, D, and G), a low level of axonemal gene transcripts remained detected in female gametocytes likely due to the transcription leaking. However, no axonemal proteins are expressed in female gametocytes [71] (Fig S11B, E, and H), suggesting a post-transcription regulation for the axonemal genes in female gametocytes. In this study, we found that enforced genomic deletion of the retained intron (*kinesin8b* intron1, *PF16* intron1, *dlc1* intron4, and *PY17X_1109100* intron1) not only restored expression of the axonemal proteins (Kinesin8b, PF16, Dlc1, and PY17X_1109100) in RBPm1-null male gametocytes (Figure 5A to L), but also unexpected resulted in low level expression of PF16, Dlc1, and PY17X_1109100 in the counterpart female gametocytes (Fig S11B, E, and H). These results confirmed the low level of these axonemal gene transcripts in female gametocytes. In addition, the translation failure of these transcripts is caused by IR in female gametocytes. In agreement with this notion, RT-PCR confirmed IR in these axonemal gene transcripts from the purified female gametocytes (Fig S11M, N, and O). As a native control, Kinesin8B was not detected in female gametocytes after deletion of the *kinesin8b intron1* (Fig S11J and K), fitting with the extremely low transcription of *kinesin8b* in female gametocytes (Fig S11L and P). Together, we proposed dual roles of RBPm1-target introns in axonemal gene expression at male and female gametocytes respectively (Fig S11Q). In male gametocytes, RBPm1 (as a key)-directed splicing of axonemal intron (as a lock) allows protein expression of the axonemal genes for axoneme assembly. In female gametocytes, dual blockage (weak transcription and intron retention) prevents protein expression of the axonemal genes.

## Materials and methods

### Animals and ethics statement

Animal experiments were conducted in accordance with approved protocols (XMULAC20190001) by the Committee for Care and Use of Laboratory Animals of Xiamen University. Female ICR mice (5-6 weeks old) were obtained from Animal Care Center of Xiamen University (at 22-24°C, relative humidity of 45–65%, a 12 h light/dark cycle) and were used for parasite propagation, drug selection, parasite cloning, and mosquito feeding. Larvae of *Anopheles stephensi* mosquitoes (*Hor* strain) were maintained under 28°C, 80% relative humidity, and a 12-h light/12-h dark condition in an insect facility. Adult mosquitoes were fed with a 10% (w/v) sucrose solution containing 0.05% 4-aminobenzoic acid and maintained at 23°C.

### Plasmid construction

The CRISPR/Cas9 plasmid pYCm was utilized for gene editing [32, 83]. To construct plasmids for gene tagging, the 5’- and 3’-flanking sequences (300 to 700 bp) at the designed insertion site of target genes were amplified as homologous templates. DNA fragments encoding 6HA, GFP, or 4Myc placed between them and in-frame with the target gene. To construct plasmids for N-terminal tagging, left homologous arms comprised 300 to 700 bp sequences upstream of the start codon, while right homologous arms comprised 300 to 700 bp sequences of the upstreaming sequences of the target gene. A DNA fragment encoding HA or 4Myc was then inserted between the left and right homologous arms in-frame with the target gene. To construct the plasmids for gene knockout, left and right homologous arms consisted of 400 to 700 bp sequences upstream and downstream of the coding sequences of the target gene. To construct plasmids for RRM or intron deletion, the RRM or intron and 200 to 700 bp sequences upstream and downstream of the RRM or intron were PCR-amplified and inserted into specific restriction sites in pYCm, followed by RRM or intron deletion using PCR-based site-directed mutagenesis with mutation primers. To construct the plasmids for inserting the intron into the *gep1* gene, the left and right homologous arms were composed of *gep1* coding sequences ranging from 300 to 600 bp upstream and downstream of the insertion site, respectively. The left homologous arm, intron, and right homologous arm were connected using overlap PCR. Subsequently, the fused fragment was inserted into specific restriction sites in pYCm. For each of the modifications mentioned above, at least two small guide RNAs (sgRNAs) were designed. Forward and reverse single sgRNA oligonucleotides were mixed, denatured at 95°C for 2 minutes, annealed at room temperature for 5 minutes, and ligated into the pYCm. To construct plasmids for the *bfp* reporter assay, the intact *bfp* reporter (717 bp) driven by the 5’-UTR (1755 bp) of the *hsp70* gene and the 3’-UTR (561 bp) of the *dhfr* gene were inserted into specific restriction sites between the left and right homologous arms of the *p230p* gene deletion plasmid [84]. The *kinesin8b* intron 1 (239 bp), *kinesin8b* intron 2 (148 bp), *PF16* intron 1 (276 bp), *dlc1* intron 4 (193 bp), PY17X_1109100 intron 1 (353 bp), and PY17X_1109100 intron 2 (272 bp) were inserted into the *bfp* reporter by overlap PCR. All primers and oligonucleotides used in plasmid construction are listed in Supplementary Table 2.

### Parasite transfection and genotyping

Parasite transfection and genotyping procedures were carried out as previously described [32, 83]. Briefly, *P. yoelii* 17XNL strain schizonts were isolated from infected mice using a Nycodenz density gradient (60% Nycodenz in PBS), and then electroporated with 5 μg plasmid using a Nucleofector 2b Device (Lonza, Germany). The transfected schizonts were immediately intravenously injected into a naïve mouse and pyrimethamine (Pyr) selection (6 mg/ml in drinking water) was applied the day following transfection. Pyr-resistant parasites were typically observed about 7 days after drug selection. Cloning of resistant parasites was performed by limiting dilution in mice, and genomic DNA was extracted from infected mouse blood for PCR genotyping using specific primers listed in Supplementary Table 2 to detect 5’ and 3’ homologous recombination events. PCR results for confirming correct genetic modification are summarized in Fig S12. All genetically modified parasite lines used in this study are listed in Supplementary Table 1.

### Negative selection with 5-fluorocytosine

To proceed with the next round of gene modification on the modified parasite clones, we initially employed negatively selection to remove pYCm plasmids. Briefly, a mouse infected with the modified parasite clones was given drinking water containing 2 mg/ml of 5-fluorocytosine (Sigma-Aldrich, cat. no. F6627) in a dark bottle. After approximately three days, the surviving parasites (most of which no longer carried pYCm plasmids, but a few still did) underwent limiting dilution and were injected into mice via the tail vein for cloning. Seven days later, blood smears were used to identify the mice that were infected with parasites (the parental generation), and these parasites were passed on to naïve mice for strain maintenance (the progeny). Then, the mice infected with the parental generation parasites were provided with drinking water containing 6 mg/ml of Pyr. If no surviving parasites were detected through blood smears under Pyr pressure, we considered this to be a parasite clone that no longer contained pYCm plasmids, and the corresponding progeny parasite clone could be used for the next round of gene modification.

### Gametocyte induction in mice

ICR mice were treated intraperitoneally with phenylhydrazine (80 µg/g body weight; cat. no. A600705-0025, Sangon Biotech) to induce hyper-reticulocytosis. At day three post-treatment, the mice were infected with 4×10^6^ asexual stage parasites via tail vein injection. The peak of gametocytaemia occurred on day three post-infection. Giemsa-stained thin blood films were used to count male and female gametocytes, and gametocytaemia was calculated as a percentage of the number of male or female gametocytes over the number of parasitized erythrocytes.

### Gametocyte purification

The procedures for gametocyte purification were carried out according to previously described methods [45]. Briefly, ICR mice were intraperitoneally treated with phenylhydrazine three days prior to parasite infection. Starting from 48 hours post-infection, the mice were orally administered 0.12 mg/d of sulfadiazine (cat. no. S8626, Sigma) for two days to eliminate asexual stage parasites. Approximately 1 ml of mouse blood containing gametocytes was collected from the orbital sinus and then suspended in 6 ml of gametocyte maintenance buffer (GMB, comprising 137 mM NaCl, 4 mM KCl, 1 mM CaCl_2_, 20 mM glucose, 20 mM HEPES, 4 mM NaHCO_3_, 0.1% BSA, pH 7.24-7.29). The 7 ml sample was layered on top of a 2 ml 48% Nycodenz/GMB cushion (consisting of 27.6% w/v Nycodenz in 5 mM Tris-HCl pH 7.2, 3 mM KCl, and 0.3 mM EDTA) in a 15 ml centrifugation tube. After centrifugation at 300 g for 30 minutes, purified gametocytes were collected from the interphase and washed twice with GMB for further use.

### Exflagellation assay

2.5 µl of mouse tail blood with induced gametocytes was mixed with 100 µl of exflagellation medium [RPMI 1640 supplemented with 100 µM xanthurenic acid (cat. no. D120804, Sigma) and 2 unit/ml heparin, pH 7.4] and incubated at 22°C for 10 minutes. For the subsequent 5 minutes, the number of ECs and total red blood cells were counted within a 1×1-mm square area of a hemocytometer under a light microscope. The number of male gametocytes and total red blood cells were also counted in a Giemsa-stained thin blood film. The exflagellation rate was calculated as the number of ECs per 100 male gametocytes. At least three biological replicates were conducted for each exflagellation assay.

### *In vitro* ookinete culture

Mouse blood with induced gametocytes was collected into heparin tubes and immediately added to the ookinete culture medium (consisting of RPMI 1640 medium supplemented with 25 mM HEPES, 10% fetal calf serum, 100 µM XA, pH 8.0) at a blood/medium volume ratio of 1:10. The samples were then incubated at 22°C for 16 hours. The number of ookinetes (both normal and abnormal morphology) per 100 female gametocytes was calculated as the ookinete conversion rate using Giemsa-stained thin blood films. At least three biological replicates were conducted for each *in vitro* ookinete culture assay.

### Parasite genetic crosses

ICR mice were treated intraperitoneally with phenylhydrazine to induce hyper-reticulocytosis. On day three post-treatment, an equal number (3×10^6^) of asexual stage parasites from two different gene knockout lines were mixed and injected via the tail vein into the phenylhydrazine pre-treated mice. After three days, blood samples were collected from the mice with induced gametocytes, and *in vitro* ookinete culture assays were performed. At least three biological replicates were conducted for each genetic cross.

### Mosquito transmission of the parasite

Approximately 100 female *Anopheles stephensi* mosquitoes were allowed to feed for 30 minutes on an anesthetized mouse infected with 4-6% gametocytaemia. To assess midgut infection, at day 7 post-feeding, approximately 30 mosquito guts were dissected, stained with 0.1% mercurochrome, and oocysts were counted under a light microscope. For counting salivary gland sporozoites, at day 14 post-feeding, salivary glands from approximately 30 mosquitoes were dissected, and gland sporozoites were counted using a hemocytometer. To assess infection by mosquito bites, at day 14 post-feeding, one anesthetized, naïve mouse was bitten by approximately 30 infected mosquitoes for 30 minutes. After 5 days, the transmission capability of parasites from mosquito to mouse was monitored in a Giemsa-stained thin blood film. All experiments were repeated three times.

### Flow cytometry analysis and sorting of male and female gametocytes

For measure DNA content in gametocytes, gametocytes purified from *DFsc7* or *DFsc7*;Δ*Rbpm1* lines were divided into two equal portions. One portion was immediately fixed with 4% paraformaldehyde in PBS, and the other portion was incubated with exflagellation medium at 22°C for 8 minutes to induce gametogenesis before fixation. The fixed cells were then stained with 4 µM Hoechst 33342 (cat. no. 62249, Thermo Fisher Scientific) at room temperature for 10 minutes, washed twice with PBS, and subjected to flow cytometry using a BD LSRFortessa instrument (BD Biosciences, San Jose, CA, USA). To exclude cell debris, Infected red blood cells were first selected by gating on forward scatter (FSC) and sideward scatter (SSC). Male gametocytes were then selected based on their GFP fluorescence intensity, and their Hoechst 33342 fluorescence intensity was analyzed. For gametocyte sorting, purified gametocytes from *DFsc7* or *DFsc7*;Δ*Rbpm1* lines were suspended in GMB at 4°C and sorted using a BD FACS AriaIII instrument (BD Biosciences, San Jose, CA, USA) based on their GFP and mCherry fluorescence intensity for male and female gametocytes, respectively. The purity of sorted gametocytes was confirmed by reanalyzing an aliquot of the sorted cells.

### Bulk RNA sequencing (RNA-seq)

Total RNA from 2×10^7^ sex-sorted gametocytes was extracted using TRIzol (cat. no. 15596026, Thermo Fisher Scientific) according to the manufacturer’s protocol. RNA integrity was assessed using the Agilent 2100 Bioanalyzer (Agilent Technologies, Palo Alto, CA, USA). mRNA was enriched using Oligo(dT) beads, fragmented into short fragments using a fragmentation buffer, and reverse transcribed into cDNA with random primers. Second-strand cDNA was synthesized using DNA polymerase Ⅰ, RNase H, dNTP, and buffer. The resulting cDNA fragments were purified with the QIAQuick PCR Purification Kit (cat. no. 28104, Qiagen), end-repaired, A-tailed, and ligated to Illumina sequencing adapters. The ligation products were size-selected using agarose gel electrophoresis, PCR amplified and sequenced using Illumina NovaSeq 6000 by Genedenovo Biotechnology Co., Ltd (Guangzhou, China).

### Differential expression analysis of bulk RNA sequencing data

The paired-end .fastq files generated from Illumina sequencer were trimmed using Trim Galore [85] (v0.6.10) (trim_galore --illumina -q 20 --paired --stringency 3 --length 25 -e 0.1 --fastqc --gzip) to remove Illumina sequencing adaptors and low quality reads. For further data clean, ribosomal RNA (rRNA) and transfer RNA (tRNA) were removed by using a genome alignment program HISAT2 [86] (v2.2.1) (hisat2 -p 12 -q --un-conc-gz). Cleaned reads (approximately 40 million reads per sample) were then mapped to reference *Plasmodium yoelii* 17X genome (PlasmoDB-62 release) using HISAT2 [86] (v2.2.1) (hisat2 -p 12 -q). The .bam files generated by HISAT2 were sorted (by position) and indexed by SAMtools [87] (v1.16.1) (samtools -sort | samtools index). Mapped reads were summarized using featureCounts [88] (v2.0.3). Gene expression analysis was performed in R [89] (v4.2.1). Gene expression values were calculated as transcripts per million (TPM) using the R package t-arae/ngscmdr (v0.1.0.181203) (https://github.com/t-arae/ngscmdr). Differential expression analysis was performed by the R package edgeR [90] (v3.40.2). Genes with fold change greater than 2 and False Discovery Rate (FDR) less than 0.05 are considered as differential expressed genes (DEGs). The volcano plot of DEGs were generated by the R package ggplot2 [91] (v3.4.2).

### Global intron retention analysis

A .gff file containing intron features was generated with a perl script from agat package and an in-house bash script, and converted into BED file by BEDOPS convert2bed (v2.4.41). DeepTools bamCoverage [94] (v3.5.1) was used to generate .bigwig files for peak visualization in Integrative Genomics Viewer (v2.16.1) [35], calculate the peak score of each exon and intron regions. RNA-seq reads aligned to different exon and intron regions were summarized by deepTools multiBamSummary [94] (v3.5.1). Low expressed genes with TPM < 30 were removed. To exclude potentially retained introns in parental samples, introns with peak scores higher than 50% of their neighboring exons in parental dataset were removed. To exclude low level retained introns in mutant samples, introns with peak scores lower than the 50% of their neighboring exons in mutant dataset were removed. The remaining introns were then normalized based on the gene expression level:

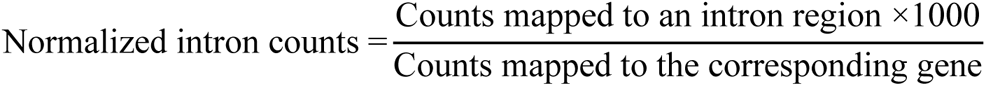

The normalized intron counts were used for differential retention analysis by the R package edgeR [90] (v3.40.2). Introns with fold change greater than 2 and FDR less than 0.05 are considered as retained introns. Retained introns were further validated manually by visualization in IGV and RT-qPCR.

### Differential expression analysis of RBP expression in male and female gametocytes

189 RBPs associated with *P. falciparum* were identified by Reddy, B.P. in 2015 [30]. Among them, 179 RBPs were found to have homologous proteins in *P. yoelii* and *P. berghei* through PlasmoDB. The differential expression of the 179 RBPs in male and female gametocytes of *P. yoelii* is based on the analysis of *P. yoelii* sex-differential gene expression in this study.

The differential expression results of the 179 RBPs in male and female gametocytes of *P. berghei* are based on the public data from Yeoh, L.M. 2017 [28]. The paired-end RNA sequencing .fastq files were downloaded from NCBI SRA database (Accession: PRJNA374918) at https://www.ncbi.nlm.nih.gov/Traces/study/?acc=PRJNA374918, and trimmed using Trim Galore [85] (v0.6.10) (trim_galore --illumina -q 20 --paired -- stringency 3 --length 25 -e 0.1 --fastqc --gzip). rRNA and tRNA were removed by HISAT2 [86] (v2.2.1) (hisat2 -p 12 -q --un-conc-gz) for data cleaning. Cleaned reads were mapped to reference *Plasmodium berghei* ANKA genome (PlasmoDB-62 release) using HISAT2 [86] (v2.2.1) (hisat2 -p 12 -q). The .bam files generated by HISAT2 were sorted (by position) and indexed by SAMtools [87] (v1.16.1). Mapped reads were summarized using featureCounts [88] (v2.0.3). Differential expression analysis was performed by the R package edgeR [90] (v3.40.2).

The differential expression results of the 189 RBPs in male and female gametocytes of *P. falciparum* are based on the public data from Lasonder E. 2016 [29]. Due to the lack of biological replicates, we turned to Cufflinks [95] (v2.2.1) for differential expression analysis in *P. falciparum*. The single-end RNA sequencing .fastq files were downloaded from NCBI SRA database (Accession: PRJNA305391) at https://www.ncbi.nlm.nih.gov/Traces/study/?acc=PRJNA305391, trimmed using Trim Galore [85] (v0.6.10) (trim_galore --illumina -q 15 --stringency 3 --length 25 -e 0.1 -- fastqc --gzip). rRNA and tRNA were removed by HISAT2 [86] (v2.2.1) (hisat2 -p 12 - q -U) for data cleaning. Cleaned reads were mapped to reference *Plasmodium falciparum* 3D7 genome (PlasmoDB-62 release) using HISAT2 [86] (v2.2.1) (hisat2 -p 12 -q --dta-cufflinks). The .bam files generated by HISAT2 were sorted (by position) and indexed by SAMtools [87] (v1.16.1). Differential expression analysis was performed by Cufflinks [95] (v2.2.1) (cuffdiff -p 8 --dispersion-method blind --library-norm-method geometric --library-type ff-firststrand).

For differential expression analysis of *P. yoelii*, *P. berghei* and *P. falciparum* RBPs, genes with fold change greater than 2 and FDR less than 0.05 are considered as DEGs. RBPs were filtered among all the genes using an in-house R script and the volcano plots of differentially expressed RBPs were generated by the R package ggplot2 [91] (v3.4.2).

### Antibodies and antiserum

The following primary antibodies were utilized: rabbit anti-HA (cat. no. 3724S, Cell Signaling Technology; IFA, 1:1000 dilution; IB, 1:1000 dilution), rabbit anti-mCherry (cat. no. ab167453, Abcam; IFA, 1:1000 dilution), rabbit anti-histone H3 antibody (cat. no. ab1791, Abcam; IFA, 1:1000 dilution), rabbit anti-Myc (cat. no. 2276S, Cell Signaling Technology; IFA, 1:1000 dilution; IB, 1:1000 dilution), mouse anti-α-Tubulin (cat. no. T6199, Sigma-Aldrich; IFA, 1:1000 dilution; IB, 1:1000 dilution; U-ExM, 1:500 dilution), mouse anti-β-Tubulin (cat. no. T5201, Sigma-Aldrich; IB, 1:1000 dilution) and mouse anti-HA (cat. no. sc-57592, Santa Cruz Biotechnology; IFA, 1:200 dilution).

The secondary antibodies employed were Alexa Fluor 555 goat anti-rabbit IgG (cat. no. A-21428, Thermo Fisher Scientific; IFA, 1:1000 dilution), Alexa Fluor 488 goat anti-rabbit IgG (cat. no. A-31566, Thermo Fisher Scientific; IFA, 1:1000 dilution), Alexa Fluor 555 goat anti-mouse IgG (cat. no. A-21422, Thermo Fisher Scientific; IFA, 1:1000 dilution; U-ExM, 1:500 dilution), Alexa Fluor 488 goat anti-mouse IgG (cat. no. A-11001, Thermo Fisher Scientific; IFA, 1:1000 dilution), Alexa Fluor 488 goat anti-mouse TER-119 (cat. no. 116215, BioLegend; IFA, 1:500 dilution), Alexa Fluor 488 conjugated streptavidin (cat. no. S32354, Invitrogen, IFA, 1:1000 dilution), HRP-conjugated goat anti-rabbit IgG (cat. no. ab6721, Abcam; IB, 1:5000 dilution) and HRP-conjugated goat anti-mouse IgG (cat. no. ab6789, Abcam; IB, 1:5000 dilution).

The antiserum, including rabbit anti-BiP (IB, 1:1000 dilution) and rabbit anti-P28 (IFA, 1:1000), were previously prepared in our laboratory.

### Live cell imaging

To perform live cell imaging of asexual blood stage, gametocyte, and salivary gland sporozoite, the *Rbpm1::gfp* parasites were first collected in 200 μl of PBS. They were stained with 4 µM Hoechst 33342 at room temperature for 10 minutes, and transferred to a 15 mm glass bottom cell culture dish (cat. no. 801002, NEST, China). Imaging was conducted using a Zeiss LSM 780 confocal microscope at a magnification of 100×. For live cell imaging of oocysts, infected midguts of *Rbpm1::gfp* were dissected in PBS on day 7 post-infection. The midguts were then stained with 4 µM Hoechst 33342 at room temperature for 10 minutes. They were mounted on a microscope slide, covered with a cover slip, and sealed with nail varnish. Imaging was conducted using a Zeiss LSM 780 confocal microscope at a magnification of 40×.

### Immunofluorescence assay

The parasites were fixed in 4% paraformaldehyde in PBS and then transferred to a 24-well cell plate with poly-L-lysine-coated coverslips. To immobilize the fixed parasites onto the glass slides, the plate was centrifuged at 550 g for 5 minutes. Next, the parasites were permeabilized with 0.1% Triton X-100 in PBS at room temperature for 10 minutes. The samples were blocked with 5% BSA/PBS at 4°C overnight, followed by incubation with primary antibodies diluted in 5% BSA/PBS at room temperature for 1 hour. After 3 washes in PBS, the samples were incubated with fluorescent conjugated secondary antibodies diluted in 5% BSA/PBS at room temperature for 1 hour. Hoechst 33342 diluted 1:5000 in PBS was added to the samples and incubated at room temperature for 15 minutes. The coverslips were washed three times in PBS, mounted in 90% glycerol, and sealed with nail varnish. Images were acquired using a Zeiss LSM 780 confocal microscope with 100× magnification.

### Ultrastructure expansion microscopy (U-ExM)

The procedures were performed as previously described in detail [96]. Purified gametocytes from 17XNL or Δ*Rbpm1* line were fixed with 4% paraformaldehyde in PBS and transferred to a 24-well cell plate with poly-D-lysine-coated coverslips. The samples were then centrifuged to sediment onto the coverslips. Subsequently, the coverslips were incubated overnight at 37°C in a mixture of 1.4% formaldehyde (cat. no. F8775, Sigma-Aldrich) and 2% acrylamide (cat. no. A4058, Sigma-Aldrich) in PBS. After that, the coverslips were gelled in a monomer solution [23% sodium acrylate (cat. no. 408220, Sigma-Aldrich), 10% acrylamide, 0.1% N,N’-methylenbisacrylamide (cat. no. M1533, Sigma-Aldrich), 1×PBS] supplemented with TEMED and APS at 37°C for 1 hour. Once the gels had completely polymerized, the coverslips (+ gels) were transferred to a 6-well plate filled with 1 ml of denaturation buffer (200 mM SDS, 200 mM NaCl, 50 mM Tris-HCl pH 8.8) and incubated at room temperature for 15 minutes to allow the gels to detach from the coverslips. The gels were then transferred into 1.5 ml Eppendorf tubes filled with denaturation buffer and incubated at 95°C for 30 minutes. After denaturation, the gels were transferred into 10 cm dishes and incubated with ddH_2_O at room temperature overnight for the first round of expansion. Subsequently, the gels were incubated with PBS for gel shrinkage, followed by transfer into a 6-well plate filled with 500 µl of mouse anti-α-Tubulin diluted in 2% BSA/PBS, and incubated at room temperature for 3 hours. The gels were washed 3 times with PBS before incubation with anti-mouse Alexa 555 diluted in 2% BSA/PBS at room temperature for 3 hours. After 3 washes in PBS, the gels were transferred into 10 cm dishes and incubated with ddH_2_O at room temperature for the second round of expansion. Subsequently, gel blocks of approximately 5×5 mm size were excised from the expanded gels and placed in the cavity well of cavity well microscope slides, covered with a coverslip, and imaged using a Zeiss LSM 980 confocal microscope.

### Protein extraction and immunoblot

Asexual blood parasites, gametocytes or ookinetes were lysed in RIPA buffer (cat. no. R0010, Solaribio) containing a protease inhibitor cocktail (cat. no. HY-K0010, MedChemExpress). After ultrasonication, the lysate was centrifuged at 14,000 *g* at 4°C for 10 minutes. The resulting supernatant was mixed with SDS-PAGE loading buffer and heated at 95°C for 5 minutes. Following SDS-PAGE separation, samples were transferred to a PVDF membrane (cat. no. IPVH00010, Millipore) and blocked with 5% milk in 1× TBST (20 mM Tris-HCl pH 7.5, 150 mM NaCl, 0.1% Tween20) at 4°C overnight. PVDF membranes were then incubated with primary antibodies at room temperature for 1 hour. After washing with 1× TBST, the membranes were incubated with an HRP-conjugated secondary antibody and then washed again with 1× TBST. Finally, the membranes were visualized using a high-sensitivity ECL chemiluminescence detection kit (cat. no. E412-01, Vazyme), and the light emission was recorded either by X-ray film or by Azure Biosystems C280 (Azure Biosystems, USA).

### Isolation of nuclear and cytoplasmic fractions

The procedures were performed with modifications according to the previous study [97]. Nycodenz-purified gametocytes were first released from red blood cells by incubating them with 0.15% saponin/PBS on ice for 5 minutes and then washed twice with ice-cold PBS. The parasite pellet was resuspended in ice-cold lysis buffer (20 mM Hepes pH 7.9, 10 mM KCl, 1.5 mM MgCl_2_, 1 mM EDTA, 1 mM EGTA, 1 mM DTT, and 0.65% Nonidet P-40) supplemented with protease inhibitor cocktail. The lysate was transferred to a 1 ml Dounce tissue grinder and homogenized gently for 80 strokes on ice. Nuclei were pelleted at 9,000 *g* at 4°C for 10 minutes, and the resulting supernatant represented cytoplasmic fractions. The nuclear pellet was washed twice with ice-cold lysis buffer before resuspension in one pellet volume of high salt buffer (20 mM Hepes pH 7.8, 1 M KCl, 1 mM EDTA, 1 mM EGTA and 1 mM DTT) supplemented with a protease inhibitor cocktail. After vigorous shaking at 4°C for 30 minutes, the extract was centrifuged at 14,000 *g* at 4°C for 10 minutes, and the resulting supernatant represented nuclear fractions. Immunoblotting was performed to analyze the proteins in each fraction.

### Protein immunoprecipitation

Nycodenz-purified gametocytes containing 3×10^7^ male gametocytes were lysed in 1 ml lysis buffer (0.01% SDS, 20 mM Tris-HCl pH 8.0, 50 mM NaCl, 1 mM DTT) supplemented with protease inhibitor cocktail. The lysate was transferred to a 1 ml Dounce tissue grinder and homogenized gently for 100 strokes on ice. The homogenate was transferred to an Eppendorf tube and incubated on ice for 10 minutes before centrifugation at 14,000 *g* at 4°C for 10 minutes. The resulting supernatant was divided into two equal portions, with one portion mixed with 20 µl pre-balanced anti-GFP nanobody agarose beads (cat. no. KTSM1301, KT HEALTH) and the other portion mixed with anti-Myc nanobody agarose beads (cat. no. KTSM1306, KT HEALTH). Both portions were incubated at 4°C for 2 hours with rotation. The beads were then washed three times with lysis buffer before elution with SDS-PAGE loading buffer, followed by incubation at 95°C for 5 minutes. Immunoblotting was performed on equal volumes of the supernatant samples.

### Transmission electron microscopy

Nycodenz-purified gametocytes were fixed at 8 mpa and 15 mpa in 2.5% glutaraldehyde in 0.1 M phosphate buffer at 4°C overnight, as previously described [98]. Then, the samples were post-fixed in 1% osmium tetroxide at 4°C for 2 hours, treated *en bloc* with uranyl acetate, dehydrated, and embedded in Spurr’s resin. Thin sections were sliced, stained with uranyl acetate and lead citrate, and examined in an HT-7800 electron microscope (Hitachi, Japan).

### TurboID-based proximity-labeling and biotinylated protein pull-down

Nycodenz-purified gametocytes containing 1×10^8^ male gametocytes from either the *Rbpm1::TurboID* or *Rbpm1::T2A::TurboID* line were incubated with 50 µM biotin (cat. no. B4639, Sigma-Aldrich) at 37°C for 20 minutes. After biotinylation, the parasites were pelleted, washed thrice with 1 ml ice-cold PBS to remove excess biotin, and then lysed with RIPA buffer containing a protease inhibitor cocktail via ultrasonication. The lysate was incubated on ice for 10 minutes before centrifugation at 14,000 *g* at 4°C for 10 minutes. The supernatant was then mixed with 50 µl pre-balanced streptavidin sepharose (cat. no. SA10004, Thermal Scientific) at 4°C overnight. The beads were washed five times with 1 ml ice-cold RIPA buffer and then washed five times with 1 ml ice-cold PBS. The washed beads were resuspended in 200 μl 100 mM Tris-HCl pH 8.5 followed by digestion with 1 µg trypsin at 37°C overnight.

### Peptide desalting and mass spectrometry

Trifluoroacetic acid (TFA; cat. no. T6508, Sigma-Aldrich) was added to the trypsin-digested sample to a final concentration of 1%, and the precipitation of sodium deoxycholate was removed by centrifugation. The resulting supernatant was desalted using in-house-made StageTips that were packed with SDB-RPS (cat. no. 2241, 3M EMPORE) and conditioned with 50 μl of 100% acetonitrile (ACN; cat. no. 34851, Sigma-Aldrich). After loading the supernatant onto the StageTips, centrifugation was performed at 3,000 g for 5 minutes. The StageTips were then washed twice with 50 μl of 1% TFA/isopropyl alcohol (cat. no. I9030, Sigma-Aldrich) followed by a wash with 50 μl of 0.2% TFA. The peptides were eluted in glass vials (cat. no. A3511040, CNW Technologies) using 80% ACN/5% NH_4_OH and dried at 45°C using a vacuum centrifuge (cat. no. 5305, Eppendorf, Hamburg, Germany). The peptide samples were resolved in 2% ACN/0.1FA for LC-MS analysis. Liquid chromatography was performed on an ultra-high pressure nano-flow chromatography system (Elute UHPLC, Bruker Daltonics). Peptides were separated on a reversed-phase column (40 cm × 75 μm i.d.) at 50°C packed with 1.8 µm 120 Å C18 material (Welch, Shanghai, China) with a pulled emitter tip. A solution is 0.1% FA in H_2_O, and B solution is 0.1% FA in ACN. The gradient time is 60 minutes and the total run time is 75 minutes including washes and equilibration. Peptides were separated with a linear gradient from 0%-5% B within 5 minutes, followed by an increase to 30% B within 55 minutes and further to 35% B within 5 minutes, followed by a washing step at 95% B and re-equilibration. LC was coupled online to a hybrid TIMS quadrupole time-of-flight mass spectrometer (Bruker timsTOF Pro) via a CaptiveSpray nano-electrospray ion source. We performed data-dependent data acquisition in PASEF mode with 10 PASEF scans per topN acquisition cycle. Singly charged precursors were excluded by their position in the m/z- ion mobility plane and precursors that reached a ‘target value’ of 20,000 a.u. were dynamically excluded for 0.4 minutes. We used 100 ms to accumulate and elute ions in the TIMS tunnel. The MS1 m/z-range was acquired from 100-1700, and the ion mobility range from 1.5 to 0.7 V cm^−2^. For data-independent acquisition, we adopted the isolation scheme of 25 Da × 32 windows to cover 400-1200 mz.

DIA files (raw) files were input to DIA-NN (v1.8.1) [99]. FASTA files downloaded from https://www.uniprot.org (UP000072874) were added. “FASTA digest for library-free search” and “Deep learning-based spectra, RTs and IMs prediction” were enabled. “Generate spectral library” was also enabled. “Protein inference” was set to “gene”. Other parameters were kept at their default settings. The protein groups and precursor lists were filtered at 1% FDR, using global q-values for protein groups and both global and run-specific q-values for precursors.

### RNA isolation, RT-PCR and RT-qPCR

Total RNA was extracted from parasites using TRIzol reagent. cDNA was synthesized with the HiScript II 1st Strand cDNA Synthesis Kit (+gDNA wiper) (cat. no. R212-02, Vazyme), using provided random hexamers, and utilized for PCR or qPCR analysis. qPCR was performed using 2×RealStar Green Fast Mixture (cat. no. A301-101, GenStar) with the following cycling program: a single incubation at 95°C for 30 seconds, followed by 40 cycles (95°C for 5 seconds, 60°C for 40 seconds) on a CFX96 Real-Time PCR System (Bio-Rad, Hercules, CA, USA). The housekeeping gene *GAPDH* (PY17X_1330200) was used as a reference gene in the RT-qPCR. The relative expression was calculated using the 2^-ΔΔCt^ method. The primers used for RT-PCRs and RT-qPCRs are listed in Supplementary Table 2.

### UV crosslinking RNA immunoprecipitation (UV-RIP)

The Nycodenz-purified gametocytes, containing 6×10^7^ male gametocytes in 6 ml ice-cold PBS, were placed in 10 cm dishes. Subsequently, they were irradiated using an HL-2000 HybriLinker (UVP, Upland, CA, USA) with 254 nm UV light at intensities of 400 mJ/cm^2^ and 200 mJ/cm^2^. The gametocytes were then collected, centrifuged, and resuspended in 1 ml lysis buffer (1% TritonX-100, 50 mM Tris-HCl pH 7.4, 150 mM NaCl, 1 mM EDTA, 1 mM EGTA) supplemented with 400 U/ml RNaseOUT (cat. no. 10777019, Thermo Fisher Scientific) and a protease inhibitor cocktail. The lysate was transferred to a 1 ml Dounce tissue grinder and gently homogenized for 100 strokes on ice. The homogenate was then transferred to a tube and incubated at 4°C for 25 minutes with rotation, followed by treatment with 30 U TURBO DNase (cat. no. AM2238, Thermo Fisher Scientific) at 37°C for 15 minutes. The lysates were centrifuged at 14,000 *g* and 4°C for 10 minutes. The supernatant was divided into two equal parts. One part was mixed with 20 µl of anti-GFP nanobody agarose beads (cat. no. KTSM1301, KT HEALTH), and the other part was mixed with 20 µl of anti-HA nanobody agarose beads (cat. no. KTSM1305, KT HEALTH). The mixtures were incubated with rotation at 4°C for 2 hours. The beads were washed six times with 500 µl RIP wash buffer (cat. no. CS203177, Millipore) at 4°C and then incubated with 117 µl RIP wash buffer, 15 µl 10% SDS and 18 µl 10 mg/ml proteinase K (cat. no. CS203218, Millipore) at 55°C for 30 minutes. RNA was isolated using phenol-chloroform extraction, and the purified RNA was reverse transcribed with random hexamer primers and determined by RT-qPCR.

### *In vitro* RNA transcription (IVT)

To prepare biotinylated probes for Figure 6O and P, IVT templates with T7 RNA polymerase promoter were obtained by PCR using the *P. yoelii* genome as a template. For Figure 7H, IVT templates with T7 RNA polymerase promoter were obtained by PCR using the plasmid used in the *bfp* reporter assay as a template. Supplementary Table 2 provides a list of primers used to obtain the IVT templates. Subsequently, Biotinylated RNA was produced using a MEGAscript kit (cat. no. AM1334, Thermo Fisher Scientific) and a biotin RNA labeling mix (cat. no. 11685597910, Roche). To create a 20 μl reaction volume, 1 μg of PCR-amplified IVT templates were incubated at 37°C for 2 hours with 2 μl of 10× reaction buffer, 2 μl of T7 RNA polymerase enzyme mix, 2 μl of biotin RNA labeling mix, and RNase-free water. The DNA templates were then removed from the RNA using TURBO DNase, and the biotinylated RNA was purified using the RNAclean Kit (cat. no. 4992728, TIANGEN). In this process, the *kinesin8b* I4 probe, *kinesin8b* I1 probe, *PF16* E1 probe, and *PF16* I1 probe all have a length of 500 nt. Additionally, the *kinesin8b* I4 probe, *kinesin8b* I1 probe, and *PF16* I1 probe span the corresponding intron sequences.

### RNA pull-down

Biotinylated RNA pull-down was performed using an RNA pulldown Kit (cat. no. Bes5102, BersinBio) following the manufacturer’s protocol. Briefly, 1 µg of biotinylated RNA was denatured at 90°C for 2 minutes and immediately cooled on ice for 2 minutes. The denatured RNA was then incubated with RNA structure buffer and RNase-free water at room temperature for 20 minutes to facilitate RNA secondary structure formation. For cell lysate preparation, Nycodenz-purified gametocytes containing 3×10^7^ male gametocytes were lysed by RIP buffer, and the resulting lysate was centrifuged at 14,000 *g* at 4°C for 10 minutes. The supernatant was then incubated with DNase Ⅰ and agarose beads to remove the chromosomes, followed by incubation with folded RNAs, streptavidin-coupled beads, and RNase inhibitor at room temperature for 2 hours. The beads were subsequently washed five times with NT2 buffer at 4°C, and proteins were retrieved from the beads by rinsing them with protein elution buffer. The retrieved proteins were then subjected to immunoblot assay.

### *bfp* reporter assay

The Nycodenz-purified gametocytes from either *DFsc7* or *DFsc7*;Δ*Rbpm1* lines, which contain a *bfp* expression cassette in the *p230p* locus, were suspended in 200 μl of GMB. The samples were then transferred to a 15 mm glass bottom cell culture dish and imaged using a Zeiss LSM 780 confocal microscope at room temperature with 100× magnification. The laser illumination was set at 561 nm (mCherry), 491 nm (GFP) and 405 nm (BFP). BFP-positive parasites indicated that the intron in the *bfp* expression cassette had been spliced.

### Other bioinformatic analysis and tools

The genomic sequences of target genes were downloaded from the PlasmoDB database (http://plasmodb.org/plasmo/). The sgRNAs of target gene were designed using EuPaGDT (http://grna.ctegd.uga.edu/). The analysis of flow cytometry data was performed using the FlowJo software (Tree Star, Ashland, OR, USA). The Gene Ontology (GO) enrichment analysis was performed using PlasmoDB. Statistical analysis was performed using GraphPad Prism (GraphPad Software Inc., San Diego, CA, USA) with either a two-tailed Student’s *t*-test or Mann-Whitney test as appropriate. Error bars represent the standard error of the mean (SEM) for triplicate experiments. *p* values were indicated in the figures above the two groups being compared, with a value < 0.05 considered significant. The protein signal on the blotting membrane was quantified using ImageJ software (NIH, Bethesda, MD, USA), and the background was subtracted from each signal. Each signal was then normalized to the Bip signal.

### Data availability

RNA-seq data for the *Plasmodium yoelii* gender-specific gametocyte transcriptome is available via the Gene Expression Omnibus database under the accession number GSE222860. RNA-seq data for the *Rbpm1* knockout transcriptome is available under accession number GSE223170. The mass spectrometry proteomic data can be accessed through ProteomeXchange with identifier PXD044094 (https://www.iprox.cn//page/project.html?id=IPX0006804000). All codes, scripts and supporting files were uploaded to GitHub at https://github.com/xiaolimo29/Intron_retention.

## Acknowledgments

This work was supported by the National Natural Science Foundation of China (31772443, 31872214, 31970387), the Natural Science Foundation of Fujian Province (2019J05010), and the 111 Project sponsored by the State Bureau of Foreign Experts and Ministry of Education of China (BP2018017).

## Author contributions

J.G. and J.Y. designed the study. J.G., P.W., X.Z., W.L., and X.Z. generated modified parasites. J.G. performed phenotype analysis, protein analysis, electron microscopy, RNA analysis, and reporter assays. P.W. conducted mass spectrometry and protein analysis. X.M. performed the bioinformatics analysis. L.J., J.L., H.C. and J.Y. supervised the work. J.G., X.M., and Y.J. wrote the manuscript.

